# Using top-down modulation to optimally balance shared versus separated task representations

**DOI:** 10.1101/2021.06.02.446735

**Authors:** Pieter Verbeke, Tom Verguts

**Author notes:** **Data availability statement:** Code to simulate the model and analyze the resulting data is provided in our GitHub repository: https://github.com/CogComNeuroSci/PieterV_public/tree/master/Gating.

## Abstract

Human adaptive behavior requires continually learning and performing a wide variety of tasks, often with very little practice. To accomplish this, it is crucial to separate neural representations of different tasks in order to avoid interference. At the same time, sharing neural representations supports generalization and allows faster learning. Therefore, a crucial challenge is to find an optimal balance between shared versus separated representations. Typically, models of human cognition employ top-down modulatory signals to separate task representations, but there exist surprisingly little systematic computational investigations of how such modulation is best implemented. We identify and systematically evaluate two crucial features of modulatory signals. First, top-down input can be processed in an additive or multiplicative manner. Second, the modulatory signals can be adaptive (learned) or non-adaptive (random). We cross these two features, resulting in four modulation networks which are tested on a variety of input datasets and tasks with different degrees of stimulus-action mapping overlap. The multiplicative adaptive modulation network outperforms all other networks in terms of accuracy. Moreover, this network develops hidden units that optimally share representations between tasks. Specifically, different than the binary approach of currently popular latent state models, it exploits partial overlap between tasks.

## 1. Introduction

Humans and other rational agents need to continually learn and perform an enormous number of complex tasks. Sometimes very similar contexts require totally different actions. For instance, while soccer and handball both require to put a ball in a goal which is guarded by a keeper and some defenders, soccer requires to manipulate the ball with the feet while handball requires to manipulate the ball with the hands. In such contexts, it is important to separate stimulus-action representations between the two tasks as much as possible in order to avoid interference. However, at other times, two different contexts nevertheless require partially similar actions. Despite the fact that tennis requires to play a ball over a low-hanging net and badminton requires to play a shuttle over a higher placed net, one can partially generalize the action of swinging the racket between the two sports. Thus, in these cases, an agent can significantly benefit from partially sharing knowledge between the two tasks.

Previous research (Baxter, 2019; Franklin & Frank, 2018; Musslick et al., 2017; Vaidya, Jones, Castillo, & Badre, 2021; Zambaldi et al., 2018) indeed illustrated that sharing task representations significantly improves learning and generalization across tasks, two hallmarks of human flexibility. However, sharing task representations in a neural network severely impacts the network’s ability to perform more than one task at the same time (i.e., to multi-task; Alon et al., 2017; Musslick et al., 2017; Musslick, Saxe, Novick, Reichman, & Cohen, 2020). Moreover, shared task representations leave a network very vulnerable to overwriting previously learned information. This problem is known as catastrophic interference (French, 1999). In contrast, a network that develops separated task representations experiences less problems in multi-tasking (Musslick et al., 2020; Tsai, Saxe, & Cox, 2016) and can continually learn without forgetting (Kirkpatrick et al., 2017; Masse, Grant, & Freedman, 2018; McClelland, McNaughton, & O’Reilly, 1995; Verbeke & Verguts, 2019). However, such networks are less able to generalize and as a consequence must learn even very similar tasks (like tennis and badminton) from scratch. In sum, there exists a trade-off between sharing and separating task representations in neural networks (Musslick et al., 2017; Musslick & Cohen, 2020; Sagiv, Musslick, Niv, & Cohen, 2020).

One popular solution to deal with this sharing-separating trade-off are compositional task representations (Fidler, Boben, & Leonardis, 2009; Franklin & Frank, 2018; Lake et al., 2014; Sugita, Tani, & Butz, 2011; Tubiana & Monasson, 2017; Yang, Joglekar, Song, Newsome, & Wang, 2019). For instance, to a first approximation, knowledge of soccer can be decomposed in two basic building blocks: the goal of the task (getting the ball past the goalkeeper) and the actions (kicking the ball). This allows the agent to generalize the goal when learning to play handball but also to avoid interference by separating the actions between both sports. Hence, a novel task can be learned quickly by recombining building blocks from previously learned tasks. Indeed, generalizing information through compositional task representations received considerable attention in several cognitive domains such as language (Irsoy & Cardie, 2014; İrsoy & Cardie, 2015; Lake et al., 2014) and sensorimotor learning (Butz, Achimova, Bilkey, & Knott, 2021; Butz, Bilkey, Humaidan, Knott, & Otte, 2019; Sugita et al., 2011). Nevertheless, it is not clear which neural network configurations could learn such compositional representations (Hupkes, Dankers, Mul, & Bruni, 2020; Lake & Baroni, 2018; Lake, Ullman, Tenenbaum, & Gershman, 2017), and what the resulting compositional representations would look like. The current work aims to build upon previous cognitive and computational work to investigate which cognitive architectures can balance shared and separated task representations in typical cognitive tasks, and what type of representations successful architectures would develop.

In cognitive science, the ability to perform one task while eliminating interference from other tasks, is known as cognitive control. Influential theoretical work (Miller & Cohen, 2001), suggests that cognitive control is implemented as a top-down modulatory signal that prioritizes relevant information processing in other processing areas. Specifically, it has been suggested that the human prefrontal cortex sends modulatory signals to more posterior processing areas; such signals excite task-relevant processing pathways and inhibit task-irrelevant processing pathways (Aben, Calderon, Van den Bussche, & Verguts, 2020). Hence, the prefrontal cortex can separate information by inhibiting all processing that might interfere with the current task. This approach has proven fruitful to explain human behavior in cognitively demanding tasks (Abrahamse, Braem, Notebaert, & Verguts, 2016; Botvinick, Braver, Barch, Carter, & Cohen, 2001; Cohen, Dunbar, & McClelland, 1990; Verbeke & Verguts, 2019). However, as noted above, a complete separation between task representations would be inefficient. Indeed, in some cases the network might benefit from information transfer between similar tasks.

To study the balance between sharing and separation, we consider the nature of top-down signals. Interestingly, there exist some crucial differences in the literature with respect to how top-down modulation is implemented. For instance, while some research treats the top-down signal as any other input signal and add all inputs together (Cohen et al., 1990), other research has treated the top-down signal as multiplicative (Masse et al., 2018; O’Reilly & Frank, 2006), which allows to effectively shut down (multiply by zero) activity for irrelevant neurons. Additionally, while in most research the modulatory signal is adapted to the needs of the current task (Botvinick et al., 2001; Cohen et al., 1990; Verguts & Notebaert, 2008), other work (Bouchacourt & Buschman, 2019; Masse et al., 2018) illustrated that also random, non-adaptive modulatory signals can be sufficient to allow optimal performance on complex tasks. Hence, random signals can often meet performance of learned signals, while requiring far less computational constraints. Moreover, because less parameters need to be learned, these random modulatory signals are often faster in learning to process novel inputs. More generally, random signals have proven to be useful in constructing powerful neural networks (Lillicrap, Cownden, Tweed, & Akerman, 2016; Maass, Natschläger, & Markram, 2002). Thus, top-down modulation signals differ in whether they are additive or multiplicative and whether they are adaptive or non-adaptive.

The current work provides a systematic investigation of different types of modulation signals in balancing the trade-off between shared and separated representations. Specifically, we propose four approaches for modulation. In a first approach, non-adaptive additive modulation (N+ network) is applied. Here, for each task, a different random top-down signal contributes to the activity patterns in an additive manner. Second, in adaptive additive modulation (A+ network), top-down input is also added to the network. However, in this approach, the top-down input is treated like any other task-processing input in the sense that top-down weights are susceptible to the same (backpropagation) learning rules as the regular task-processing weights. Third, in non-adaptive multiplicative modulation (Nx network), the network inhibits and/or excites a random proportion of pathways in every task context by multiplying activation with zero (inhibition) or a random positive value (excitation). Fourth, in adaptive multiplicative modulation (Ax network), the network learns which processing pathways to excite or inhibit.

Since previous work illustrated that the impact of shared representations depends on the nature of the task environment (Musslick et al., 2017), networks are tested on three different types of input (discrete low-dimensional, continuous low-dimensional, and continuous high-dimensional; see also Figure 1). For each input type, we consider a number of tasks that differ in the amount of overlap of their stimulus-action mappings. Interestingly, for artificial agents, there is more catastrophic interference when trained in a blocked fashion. In contrast, blockwise training appears beneficial for human agents (Flesch, Balaguer, Dekker, Nili, & Summerfield, 2018). To evaluate each network’s ability to overcome interference, we thus trained our artificial networks in a blocked fashion. In sum, we test the four modulation signals on a task that requires them to optimally balance the transfer (sharing) and avoidance of interference (separating) between tasks. Network performance is evaluated in terms of accuracy and the ability to find the optimal amount of sharing between task representations.

**Figure 1.**
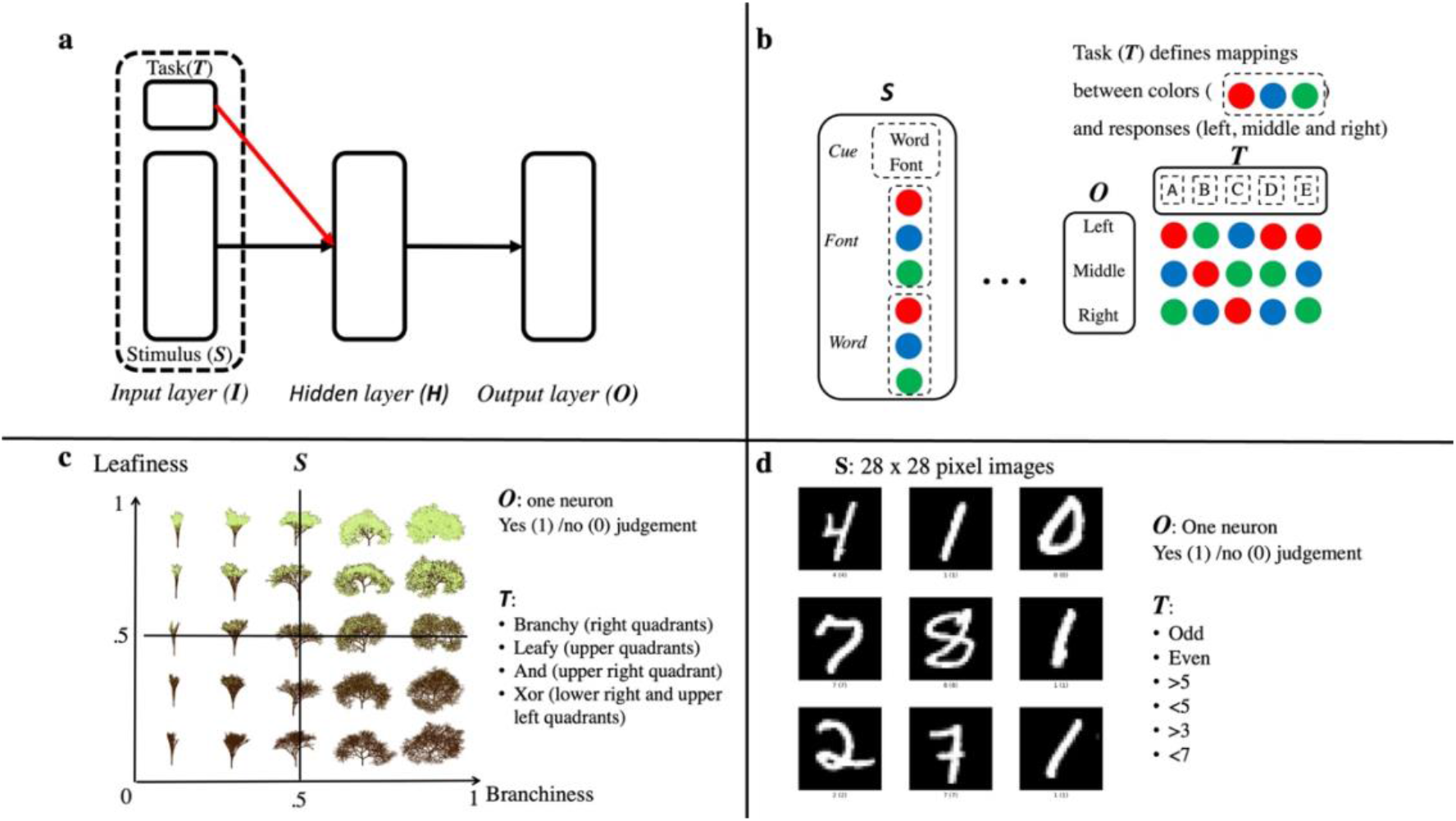
Network architecture and simulations. *a: General network architecture.* The network consists of three layers. Information flows in a feedforward manner from Input to Hidden to Output layer. The Input layer is divided in a Stimulus and Task group. The Task group sends a modulation signal (red arrow). We evaluate four different types of modulation and test the network on three types of stimulus datasets (b-d). *b: Stroop (discrete) dataset.* Stimuli are a combination of a cue, a font (color) and a (color) word. The cue indicates which other dimension (word or font) is relevant for responding. Five tasks are defined in which the mapping between three response options and three values for color (in font and word) are changed. Mappings are represented in the panel. *c: Trees (continuous low-dimensional) dataset.* The stimulus figure is adopted from Flesch et al. (2018). Current work has coded this dataset with two neurons that can take on any value between 0 and 1. Four tasks are defined in which the network needs to give a yes or no judgement (one response neuron). *d: MNIST (continuous high-dimensional) dataset.* Here, stimuli are 28 x 28 pixel images of handwritten digits from 0 to 9. Examples are taken from https://www.tensorflow.org/datasets/catalog/mnist. We defined six possible tasks in which again the network needs to give a yes/no judgement.

## 2. Methods

### 2.1 The network

Our network (Figure 1a) consists of an Input, Hidden, and Output layer. Information flows in a feedforward manner from Input to Hidden to Output layer. All neurons in each layer are fully connected to all neurons in the next layer. Neurons in the Input layer are divided in a Stimulus group and a Task group. Activation at the Hidden layer is a combination of input from the Stimulus group and a modulatory signal from the Task group. In analogy to previous work, we use

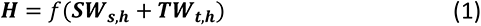

for additive modulation (e.g., Cohen et al., 1990) and

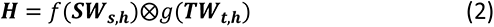

for multiplicative modulation (e.g., Masse et al., 2018). However, see the Supplementary materials for additional simulations with other implementations of modulation. In these equations, ***H, S*** and ***T*** are vectors representing activation in the Hidden, Stimulus and Task group respectively. A weight matrix between layers is represented by ***W***. The symbol ⊗ represents elementwise multiplication. The functions *f*() and *g*() represent (elementwise) nonlinear activation functions. In the main text we consider simulations in which *f*() represents a sigmoid activation function:

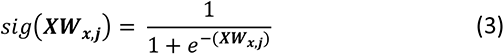

and *g*() represents a RELU activation function:

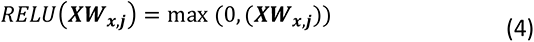

In these equations, ***X*** is a (row) vector representing activity in the sending layer and ***Wx,j*** represents a weight matrix between the sending (*x*) and receiving (*j*) layer. Thus, Hidden neurons with a negative weight from the active Task neuron are gated out (activation multiplied by 0), while activation of Hidden neurons with a positive weight, are multiplied by a positive value. The Supplementary materials present additional simulations in which different combinations of sigmoid and RELU functions are explored. Activation at the Output layer (***O***) simply follows ***O*** = *sig*(***HWh,o***). After each trial, weights are adapted by the backpropagation learning rule (Rumelhart, Hinton, & Williams, 1986):

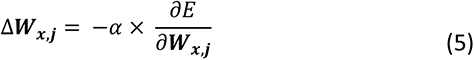

in which again *x* represents the sending layer and *j* represents the receiving layer. The parameter α > 0 represents a learning rate, and 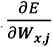 is the (partial) derivative of the error (*E*) with respect to the weights (***W***).

### 2.2 Datasets

The network was tested on three datasets (Figure 1b-d), allowing us to evaluate several combinations of input and task type. Note that the size of the Input, Output and Task layers were adapted depending on the input dataset. The first dataset had discrete (binary) low-dimensional input patterns. Specifically, we consider the classic cognitive control Stroop task (Stroop, 1935). See Figure 1b for an illustration of the network specifics for this dataset. Here, on each trial a (color) word is presented (“red”, “green” or “blue”) in a particular font (color) (red, green or blue). Additionally, a cue is provided telling the agent to respond either to the word or to the font dimension. The task consists in learning mappings between colors (red, green and blue) and response buttons (left, middle, right). Crucially, both stimulus dimensions can provide congruent evidence (e.g. “red” presented in red) or incongruent evidence (“red” presented in blue). In the latter case, the correct response depends on the cue dimension (respond to red when cue is word and to blue when cue is font). In terms of the network, we consider 8 input neurons (2 cues, 3 words and 3 colors; see also Figure 1b). Here, each stimulus consists of the activation (input value = 1) of 3 (a cue, a word and a font) out of 8 Stimulus neurons, resulting in 18 (2 instructions × 3 fonts × 3 words) possible stimuli. Additionally, we activate one Task neuron on every trial, which determines the appropriate mappings between stimuli and responses. In this task, the Output layer consists of 3 neurons. On each trial, the Output neuron with the highest activation (argmax(***O***)) is considered to be the network response. Depending on the task, each color value (red, green or blue) was mapped to one of the neurons in the Output layer. Specifically, we define five tasks (see also Figure 1b). Here, tasks A, B and C share no stimulus-action mappings. Task D represents a mix of tasks A, B and C. Specifically, D shares exactly 1/3 of stimulus-action mappings with all three other tasks. The last task E shared all stimulus-action mappings with A but activated a different neuron in the Task group (in a sense, A and E are synonyms). Note that we call this a Stroop task because the input consists of font and word input where one dimension was relevant; we did not mimic the imbalance between color naming and word reading that appears in typical Stroop tasks.

The second dataset is the Trees dataset (see also Flesch et al., 2018). Specifics for this dataset are presented in Figure 1c. In the Trees dataset, there are two Stimulus neurons which can take on any value in a range of 0 to 1 (continuous low-dimensional input). One Stimulus neuron represents the ‘leafiness’ of a tree and the other Stimulus neuron represents the ‘branchiness’ of the tree. The Output layer contained only one neuron. For this dataset, the network has to make a yes (output = 1) or no (output = 0) judgement. We defined four different tasks for this input type. One task was to respond yes to leafy trees (leafiness >.5), a second task was to respond to branchy trees (branchiness >.5), a third task required the network to respond to trees that were both leafy and branchy (AND task), and the fourth task consisted of responding to trees that were either leafy or branchy (but not both; XOR task). Note that for this dataset there were no completely (100%) dissimilar tasks. The Leafy and Branchy task share 50% of stimulus-action mappings with each other but also with the AND and XOR taks. The AND and XOR task share 25% of mappings with each other.

The third dataset consisted of images (continuous, high-dimensional input). More specifically, we used the MNIST data set (LeCun, Cortes, & Burges, 2010) which contains grey-scaled images (28 x 28 pixels) of handwritten digits from 0 to 9. Again, the Output layer consisted of one neuron. For this input type, 6 different tasks were provided. One task was to respond to odd digits (i.e., output = 1 for odd digits; output = 0 for even digits); another task required a response to even digits. A third and fourth task required the network to respond to digits that were respectively larger or smaller than 5. The fifth task consisted of responding to digits larger than 3 and the sixth task was to respond to digits smaller than 7. This resulted in a complex pattern of overlap between the different tasks. Tasks vary from 100% dissimilar (odd and even), to only 20% dissimilar (>3 and >5; <5 and <7).

### 2.3 Simulations

As described before, four versions of the network were simulated. Activation at the Hidden layer follows Equation (1) for additive modulation networks (N+ and A+), and Equation (2) for multiplicative modulation networks (Nx and Ax). In adaptive modulation networks (A+ and Ax), the weights between the Task group and Hidden layer are learned by the backpropagation rule (Equation (5)), just like the other weights. In non-adaptive modulation networks (N+ and Nx), the weights between the Task group and Hidden layer are fixed at their initial (random) values. All weights are initialized with a random value drawn from the normal distribution N(0, 1). Only for the Ax network, modulating weights (between Task and Hidden layer) had an initial random value drawn from the uniform distribution U(0, 1), such that RELU(***T***) > 0 (all gates open) at the first trial. This set up provides the most optimal initialization for each network. We illustrate network performance with other weight initialization distributions in the Supplementary materials.

All four versions of the network were tested on all three data sets. Additionally, we explored different learning rates (α) and shapes of Hidden layer. Also the shapes of the Input, Output and Task layers were adapted depending on the input dataset. For the Stroop and Trees input datasets, α took on 6 values ranging from 0 to 1 in steps of .2. We explored the network with one Hidden layer of either 12 or 24 neurons. For the MNIST dataset we used lower learning rates. Here, α took on 6 values ranging from 0 to .1 in steps of .02. For this data set, we explored performance with one Hidden layer of 400 neurons; and also with two Hidden layers (300 and 100 neurons respectively) and three Hidden layers (200, 100 and 100 neurons respectively). Note that for this dataset, the total number of Hidden neurons did not differ between architectures. Activation at the first Hidden layer (***H***1) followed Equation (1) or (2) for additive or multiplicative networks respectively. In standard simulations, activation at the second and third Hidden layer followed: ***H***i = *sig*(***H***i-1***W****Hi-1,Hi*), in which *i* is the index of the Hidden layer. Hence, the Task modulation signal was not sent directly to the deeper Hidden layer(s). However, for completeness we also explored network performance (with two hidden layers) when the Task signal was sent to only the second hidden layer, to both hidden layers or to none of the hidden layers. Results of these simulations are presented in section 3.5.

For every combination of α and shape of the Hidden layer, 25 simulations (*N* = 25) were performed for each dataset. For each simulation, 1200 or 12000 inputs were randomly sampled for the Trees and MNIST datasets respectively. Since there were only 18 stimuli (input patterns) for the Stroop dataset, we chose to repeat these 18 stimuli 75 times in each simulation, resulting in 1350 trials. Additionally, we randomly shuffled the order of tasks before a simulation. In a next step, we divided the sampled input patterns over 3 repetitions (450, 400 or 4000 trials per block for the Stroop, Trees and MNIST dataset respectively). In every repetition, the network was trained (training phase) blockwise on each task, using the predetermined input sample and order of tasks. Thus, each task was repeated 3 times in a blocked fashion. At the end of the third block, weights were frozen, and the network was tested (test phase). For this test phase a new order of tasks was generated. Each task was tested for one block of trials. For this purpose, 100 or 500 new inputs were randomly sampled from the Trees and MNIST datasets respectively. For the Stroop dataset, no new inputs could be generated so we repeated the 18 possible inputs 5 times, resulting in 90 trials.

### 2.4 Analyses

#### 2.4.1 Accuracy

To investigate whether networks suffered from catastrophic interference during learning, we computed accuracy for each task repetition (averaged over all tasks). Networks that suffer from catastrophic interference would need to relearn a task on every repetition because they would learn other tasks in between. Hence, such a network would not improve over task repetitions.

Next, we investigated the network’s ability to balance separating representations with sharing representations. More specifically, we computed accuracy for each task during the test phase. For this analysis we mainly focus on the Stroop dataset but we present results for the other datasets as well. The Stroop dataset is optimally suited for this analysis since there is a larger variation in (dis)similarities between tasks (see also the objective dissimilarity table in Figure 2) than is the case for the other datasets. More specifically, five tasks were proposed for the Stroop dataset. As described in section 2.2, three of them (A, B and C) did not share any stimulus-action mappings and thus can be totally separated. Tasks A and E however, share all stimulus-action mappings and can be fully shared. Additionally, task D shares 1/3 of its stimulus-action mappings with all other tasks. On the one hand, a full sharing of task representations would allow the network to exploit the shared mappings between A, D and E but lead to catastrophic interference in tasks B and C, illustrated by a strong decrease of accuracy in tasks B and C compared to A, D and E. On the other hand, a full separation of tasks would improve accuracy of tasks B and C (by less interference), but would also eliminate the advantage of the full overlap between A and E and the partial overlap between D and the other tasks. Hence, accuracy would be the same for all tasks. Importantly, when the network is only able to fully share or separate information it would benefit from the full overlap between A and E but would not benefit from the partial overlap between task D and the other tasks. In sum, an optimal network would find a balance between sharing and separating, resulting in an improved accuracy for tasks A, D and E while minimizing the dip in accuracy for tasks B and C.

**Figure 2.**
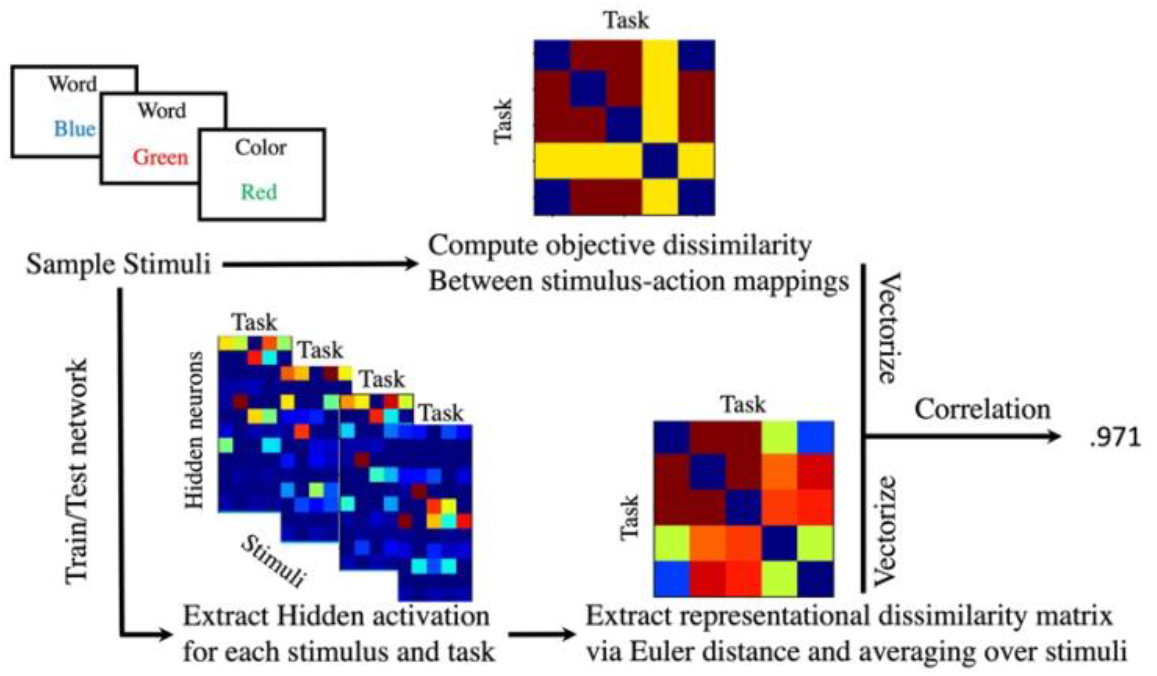
Methods. Illustration of the different steps in the representational dissimilarity analyses. Examples are shown for one simulation of the Stroop dataset with a learning rate of .6 and 12 Hidden neurons.

To evaluate overall performance of the networks we also computed accuracy during the test phase for all learning rates and all tasks.

#### 2.4.2 Representational dissimilarity

In order to analyze to what extent the networks shared or separated stimulus-action mappings across tasks, we computed how dissimilarity between tasks in terms of stimulus-action mappings was represented in the network. This analysis considers several steps. An overview of these steps in the context of the Stroop dataset is provided in Figure 2.

In a first step, we computed for each simulation the objective dissimilarity between stimulus-action mappings across tasks. Specifically, we computed a matrix where rows and columns represent the tasks, and each cell contains the proportion of stimuli that were matched with a different action across the two respective tasks (row and column).

A second step was to compute the representational dissimilarity within the network. For this purpose, we first computed the mean activation at Hidden layer for each stimulus (18 Stroop stimuli, 4 quadrants of branchy-leafy space and 10 digits in MNIST dataset) in each task across trials. Then the difference between task representations was extracted by computing the mean Euclidean distance for two tasks *T*1 and *T*2. Here,

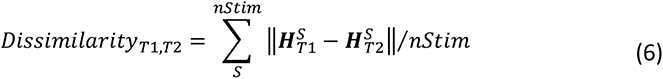

in which 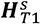 and 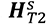 are vectors of length *nHidden*, representing the average activity for all Hidden neurons when stimulus *S* was presented to the network. Hence, we compute the Euclidean distance (indicated by÷÷-÷÷) for each stimulus (*S*) and each task pair (*T1*, *T2*). This distance is then averaged over all possible stimuli (*nStim*) to obtain one dissimilarity matrix of Hidden representations between tasks.

In a third and last step we compared the objective dissimilarity to the representational dissimilarity. Specifically, we reshaped both matrices to vectors and computed the Spearman rank correlation coefficient between these vectors. This resulted in one value of the dissimilarity correlation between objective task dissimilarity and a network’s representational task dissimilarity.

#### 2.4.3 Neural activation analyses

We performed two additional analyses to gain more insight into how the different modulatory signals organize Hidden layer activity. For this purpose, we again computed for each task the mean activation at Hidden layer for each stimulus, resulting in a matrix with size (*nStim*, *nTask*, *nHidden*). First, we investigated the distribution of activation for all stimuli and tasks across the Hidden neurons. Second, in order to visualize the network representations for each task, we reduced Hidden layer dimensionality via principal component analysis. For this purpose, we entered the activation matrix with size (*nStim*, *nTask*, *nHidden*) into the principal component analysis and approximated it by a matrix of size (*nStim*, *nTask*, 2). As a result, we could plot the representation of each stimulus in each task in a two-dimensional space.

## 3. Results

### 3.1 Accuracy

First, we evaluated the networks’ ability to separate task representations. For this purpose, we investigated whether average accuracy (during the training phase) increased over task repetitions. Networks that do not separate task sets suffer from catastrophic interference because they overwrite mappings of one task by the mappings of another task. As a result, such networks need to relearn the original task when it is presented again, and do not show any improvement over task repetitions. In Figure 3, it is observed that accuracy hardly improves for the additive networks (A+ and N+). Thus, these networks severely suffered from catastrophic interference. In contrast, for multiplicative modulation networks, in particular for the Ax network, there was a significant improvement over task repetitions. Thus, multiplicative modulation seems more efficient in separating task representations, rendering them less vulnerable to catastrophic interference during learning.

**Figure 3.**
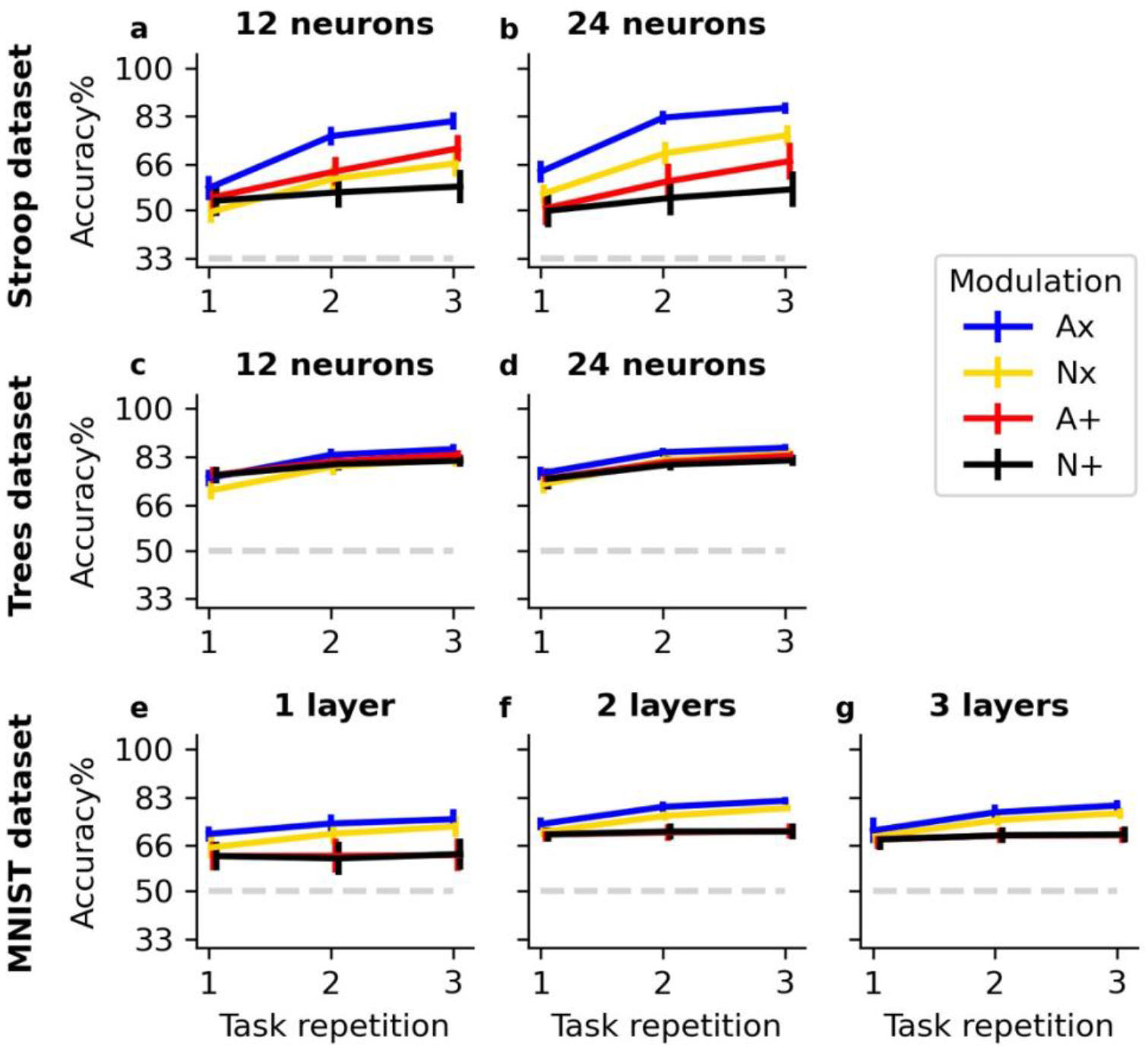
Accuracy in training phase per task repetition. Lines illustrate mean accuracy for each task repetition during the training phase averaged across all learning rates (α), all tasks and all simulations. Bars indicate 95% confidence intervals over 25 simulations. The dashed lightgrey line indicates chance level accuracy. Results are shown for different datasets (rows) and different shapes of Hidden layer (columns).

Second, we zoomed in on accuracy for each task during the test phase. Here, we focus mainly on the Stroop dataset (see section 2.2) because the Stroop dataset has a broader range of dissimilarities between tasks. Specifically, tasks A, B and C have completely dissimilar mappings, task D has a partial overlap of 1/3 with all other tasks and task E shares all mappings with task A. An optimal network would find a balance between sharing and separating, resulting in an improved accuracy for tasks A, D and E while minimizing the dip in accuracy for tasks B and C (see also section 2.4.1). In Figure 4a,b we observe a strong dip in accuracy for tasks B and C when the modulation signal was additive. This suggests that additive modulating signals are well suited for sharing task representations, but less so for separating task representations. This dip in accuracy for tasks B and C is less strong for the multiplicative modulation networks. Importantly, the Nx network has an approximately equal accuracy for all tasks (Figure 4a,b). This suggests that the Nx network has strongly separated task representations, which did not allow that network to benefit from overlap between task mappings across the different tasks. The Ax network is clearly the network that was able to optimally balance the separation and sharing of task representations, showing an advantage in accuracy for all tasks compared to the other networks, and only a small dip for tasks A and B. Note that although task D only had a partial overlap with other tasks, accuracy is equally high as for tasks A and E which fully overlapped. Hence, the Ax network does not treat sharing or separation as an all-or-none process, but also captures partial overlap.

**Figure 4.**
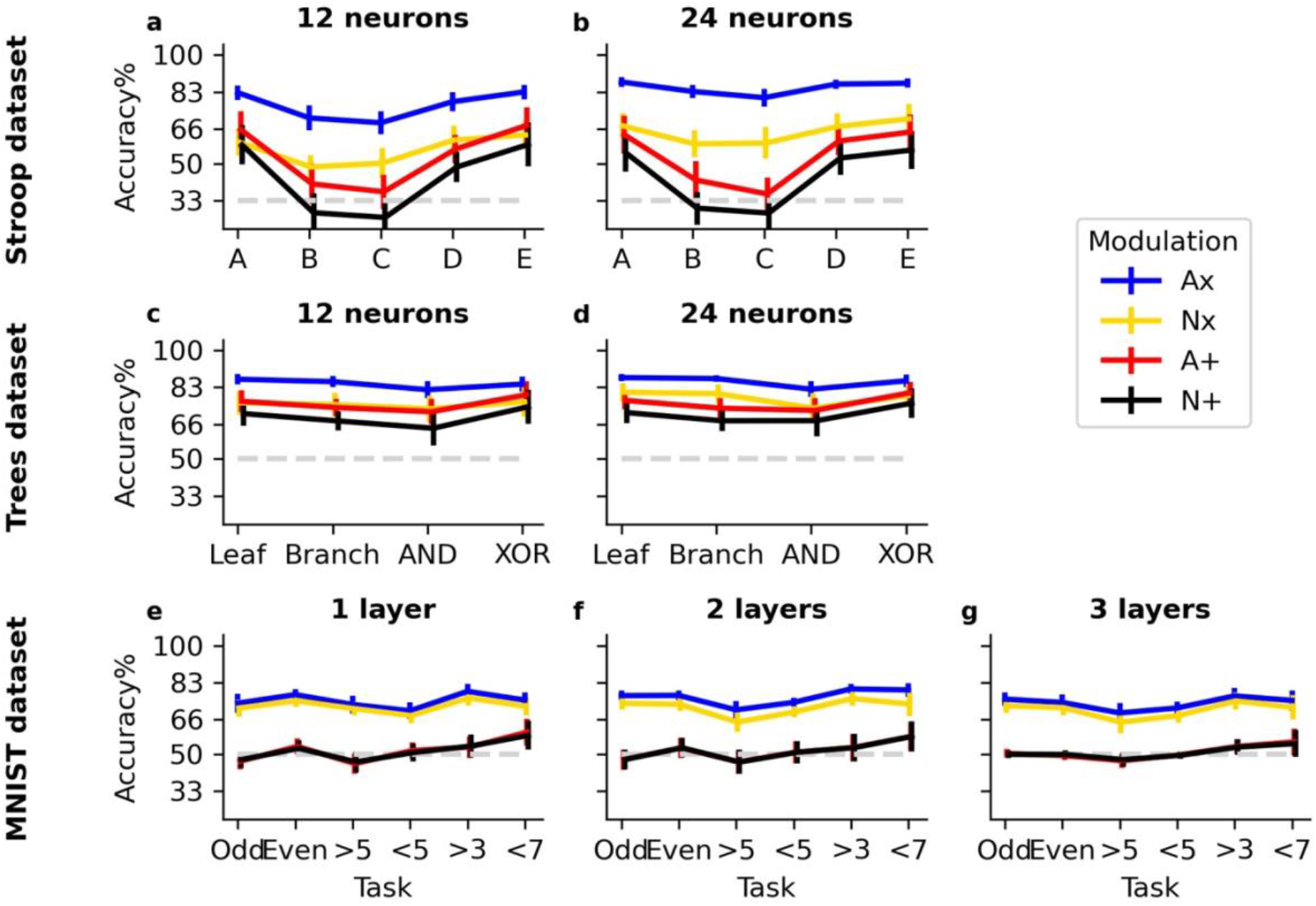
Accuracy in test phase per task. Lines illustrate mean accuracy during the test phase for each task, averaged across all learning rates (α) and all simulations. Bars indicate 95% confidence intervals over 25 simulations. The dashed lightgrey line indicates chance level accuracy. Results are shown for different datasets (rows) and different shapes of Hidden layer (columns).

We also show results for the other datasets but these are less informative in this respect, because there is no strong variability in objective task dissimilarity (see section 2.2). For the Trees dataset (Figure 4 c,d) there was an average objective dissimilarity of 50% for the leafy and branchy task, and an average dissimilarity of 41.5% for the AND and XOR tasks. For the MNIST dataset (Figure 4e-g) there was a strong variability of dissimilarity between tasks themselves (20-100%), but when averaged over all tasks, the range of dissimilarity between one task and all other tasks was rather small (between 56 and 64%). Despite the absence of variability in overall dissimilarity between tasks, it is also clear for the Trees (Figure 4c, d) and MNIST (Figure 4e, f) dataset that the Ax network performs best.

This suggestion that the Ax network is best able to find a balance between sharing and separating task representations, is supported by the fact that this network reaches a higher accuracy overall when we analyze accuracy over all tasks during the test phase. In Figure 5, it is observed that the Ax network outperforms all other networks for all learning rates, all shapes of Hidden layer and all datasets. The non-adaptive additive (N+) network performs worst. The non-adaptive multiplicative modulation (Nx) network and the adaptive additive (A+) network perform in between these other networks. Here, the Nx modulation network seems to obtain an advantage over the A+ network when there are more Hidden neurons (Figure 5b,d) and for high-dimensional (MNIST; Figure 5e,f) input datasets. In sum, multiplicative modulation outperforms additive modulation, and adaptive modulation outperforms non-adaptive modulation.

**Figure 5.**
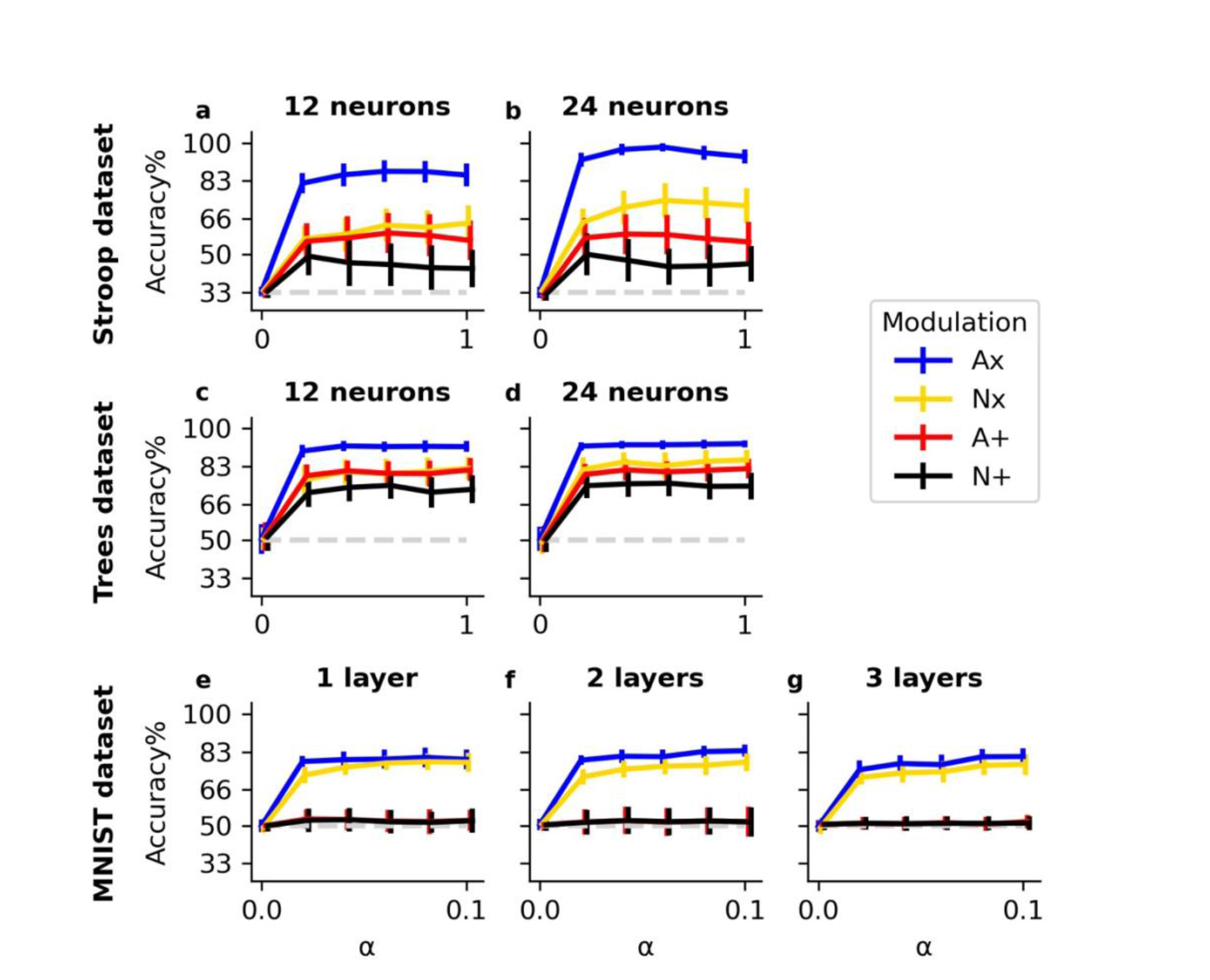
Accuracy in test phase per learning rate (α). Lines illustrate mean accuracy during the test phase for each value of α, averaged across all tasks and all simulations. Bars indicate 95% confidence intervals over 25 simulations. The dashed lightgrey line indicates chance level accuracy. Results are shown for different datasets (rows) and different shapes of Hidden layer (columns).

### 3.2 Representational dissimilarity

We next investigate whether objective dissimilarity of stimulus-action mappings between tasks was represented in the network. For this purpose, we computed for each network simulation a representational dissimilarity matrix and correlated this matrix elementwise with an objective dissimilarity matrix of stimulus-action mappings between tasks (see section 2.4.2 and Figure 2 for details). Results of this analysis are shown in Figure 6. As was already suggested by the accuracy results (section 3.1), the Ax network was clearly better at extracting the objective dissimilarity between tasks. Interestingly, the A+ network was also (although less strongly) able to capture the objective overlap between tasks for the Stroop and Trees input datasets (Figure 6a-d), but not for the MNIST dataset (Figure 6e-g).

**Figure 6.**
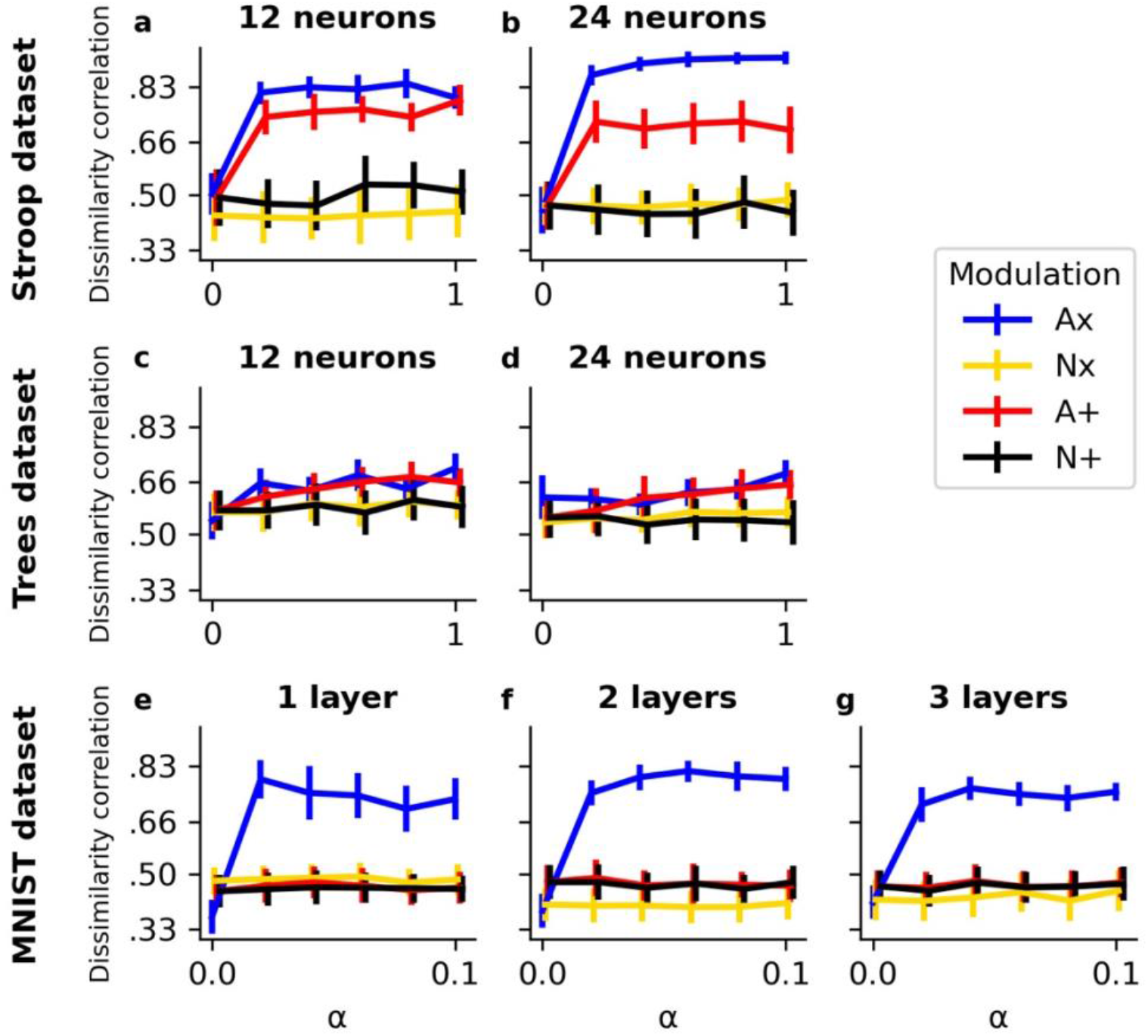
Correlation of objective and representational dissimilarity between tasks. Lines illustrate the mean task dissimilarity correlation for each value of α across all simulations. Bars indicate 95% confidence intervals over 25 simulations. Results are shown for different datasets (rows) and different shapes of Hidden layer (columns).

For the MNIST dataset, we simulated a network with one Hidden layer (Figure 6e), one with 2 Hidden layers (Figure 6f) and one with three Hidden layers (Figure 6g). Overall, there does not seem to be a strong benefit for dividing the (same number of) Hidden neurons over multiple layers in the current setup. However, only the first Hidden layer received a modulation signal. In section 3.5 we present results for simulations that modulated deeper and/or more hidden layers.

### 3.3 Neural activation analysis

To provide additional insight into how the different modulation signals organize Hidden layer activity, Figure 7 shows the distribution of activation at the Hidden layer for all networks and datasets. This is shown for the networks with 12 Hidden neurons (Stroop and Trees dataset) or 1 Hidden layer (MNIST dataset). Here, it is observed that activation distributions are strongly bimodal with peaks around 0 and 1. Note that, because of the RELU modulatory signal (see Equations (2) and (4)), activation in the multiplicative modulation networks were theoretically not bound to 1. Nevertheless, also these multiplicative modulation networks show a clear activation bound of 1 after learning. Interestingly, the multiplicative modulation networks illustrate a strong asymmetrical distribution with a higher peak of activity around zero. Especially the Nx network has a high zero-centered peak. This suggests that the Nx network is learning more sparse representations and potentially creates different groups or modules of neurons where each module learns (part of) one task. Hence, in line with what was described before, the Nx network is well suited for separating task representations. The activity distribution of the additive networks is clearly more symmetrical with a higher number of neurons that exhibit strong activity for each stimulus and task. As a result, the additive networks will probably share more neurons for representing stimuli and/or tasks. In line with what we described before, the Ax network illustrates a mixture between the properties of the Nx and additive networks and is therefore optimally suited to balance shared and separated representations.

**Figure 7.**
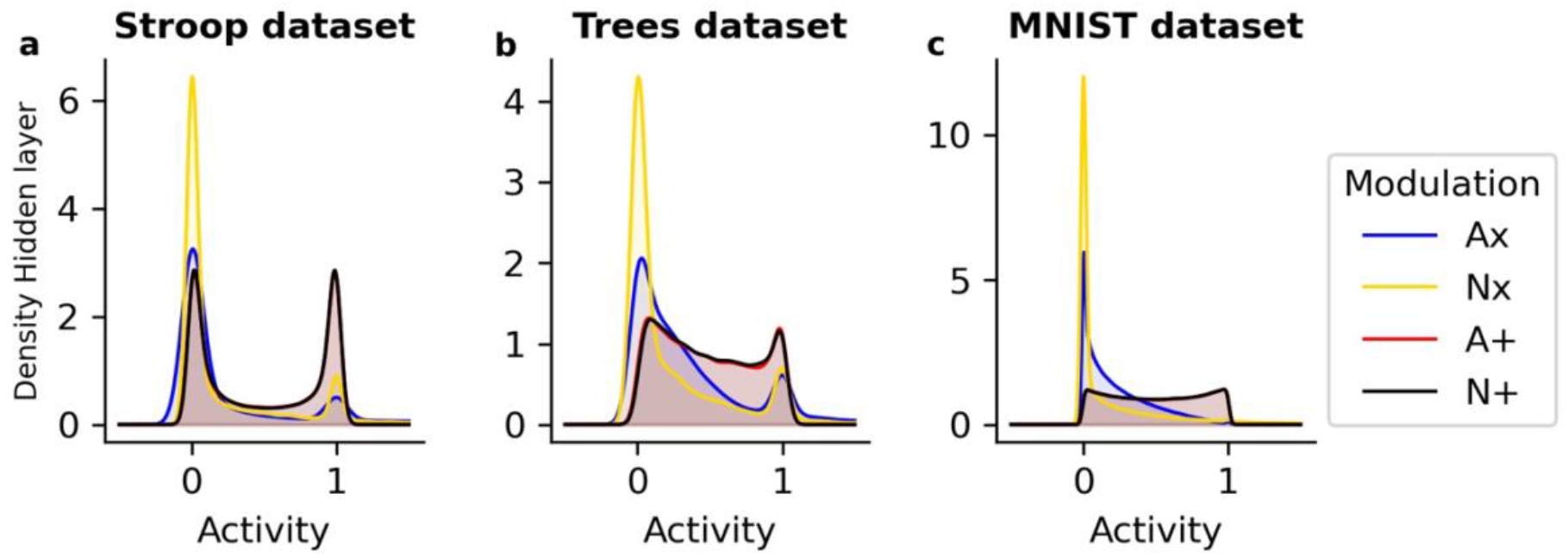
Distribution of activity at Hidden layer. The distribution of activation at the Hidden layer of the different modulation networks is shown for each dataset.

For the next analyses, we reduced Hidden layer dimensionality into two principal components with the highest eigenvalues. For the Stroop dataset these components explained on average over all simulations, networks, learning rates and shapes of Hidden layer, 54.34% of the variance (SD = 6.41%). For the Trees dataset, two components explained 71.29% of the variance (SD = 8.16%) and for the MNIST dataset, two components explained 44.95% of the variance (SD = 4.68%). In Figure 8, we show neural representations in the Hidden layer for each stimulus (dots) and each task (colors) on 2 dimensions. Notice that this analysis does not allow us to average over simulations. Hence, results are shown for one representative simulation of the network with an intermediate learning rate of α = .6 for the Stroop and Trees dataset and α = .06 for the MNIST dataset.

**Figure 8.**
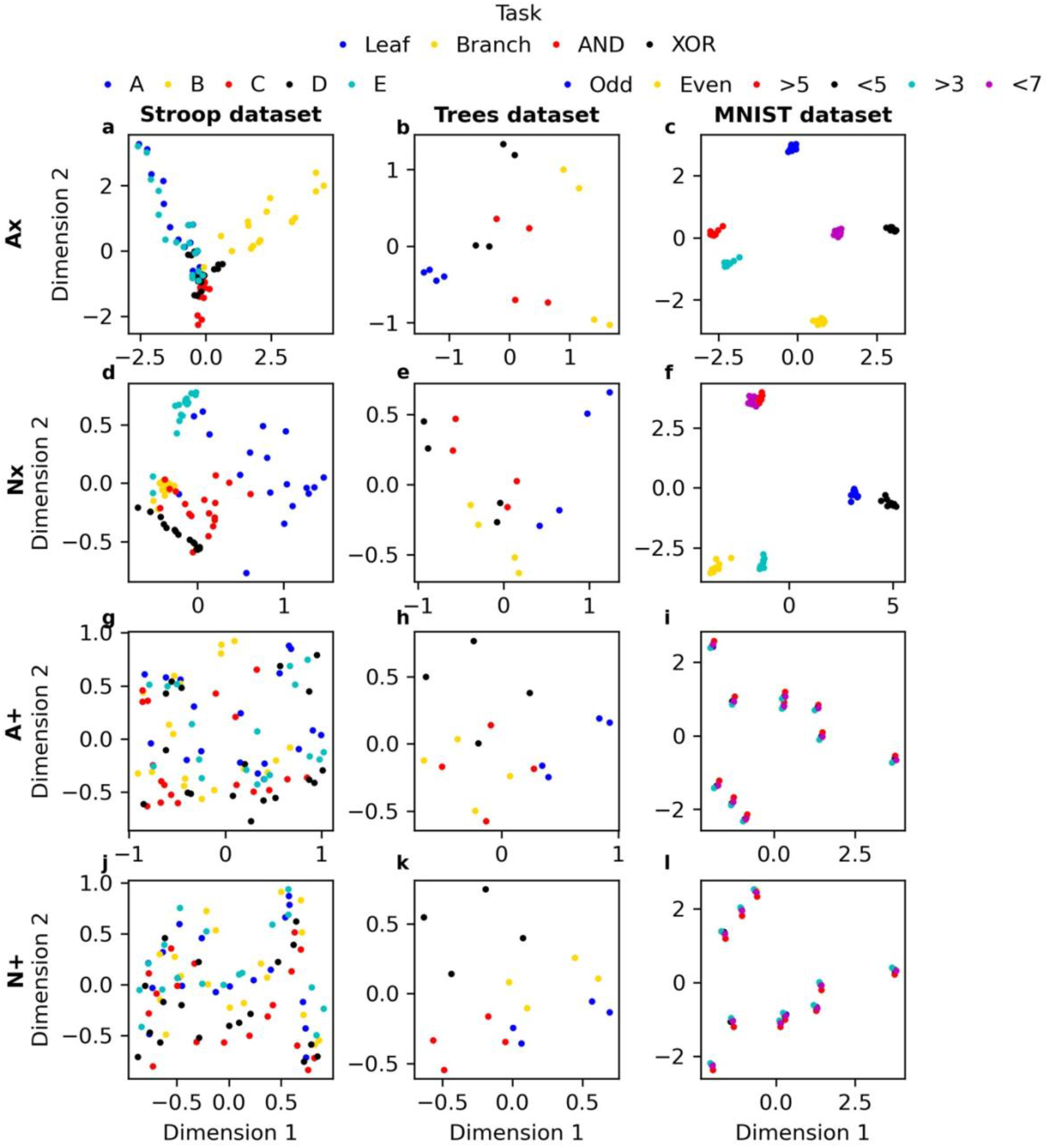
Task representations after principal component analysis. The neural representation for each stimulus (dots) and each task (colors) are shown along the first two principal components. This is shown for a representative simulation and an intermediate learning rate, for all modulation networks (columns).

Generally, results are in line with our previous findings in accuracy and representational dissimilarity analyses. For the Stroop dataset, we observe that the additive modulation networks show a tendency to share neural representations across tasks (Figure 8g,j). In contrast, the Nx network (Figure 8d) effectively separates the different tasks but fails to share tasks A and E (blue and green dots) which have no dissimilarities in their stimulus-action mappings. The Ax network however (Figure 8a), shows a remarkable ability in discovering the overall relational structure between tasks. The network finds three orthogonal axes for the three orthogonal tasks A, B and C. Task D which shares stimulus-action mappings with all previous tasks, is placed in between (at the origin of the three axes) the representations of A, B and C. Additionally, the network discovered that task E fully overlaps with task A.

For the Trees dataset (Figure 8b,e,h,k), all networks were able to separate task representations. Thus, in contrast to the Stroop input datasets, the additive networks were able to separate task representations for the Trees dataset. This explains why the difference in accuracy between networks was much smaller for the Trees dataset in comparison to other datasets (Figure 5). Note that in this dataset the inputs were significantly less complex (only 2 dimensions) than for the other datasets (18 or 784 (28^2^) dimensions for the Stroop and MNIST dataset respectively).

For the MNIST dataset, the additive networks again fail to separate task representations (Figure 8i,l). Notice that the networks extracted separate representations for the 10 digits (0-9) but shared the digit representations across all tasks. The Nx network (Figure 8f) was able to separate task representations but did not extract a clear relational structure. Again, only the Ax network (Figure 8c) was able to extract the full relational structure of the tasks. Here, two dimensions were extracted by the network. One dimension (Dimension 2) was used to separate the odd from the even tasks, the other dimension (Dimension 1) was used to separate the larger than (>5 and >3) tasks from the smaller than (<5 and <7) tasks.

### 3.4. Multilayer networks

To provide more insight into the deeper (more than one Hidden layer) networks, we provide results of the representational dissimilarity analyses in each layer separately (Figure 9a-e). This is shown for the two and three Hidden layer networks that were tested on the MNIST dataset. Remarkably, while the modulation signal is only delivered to the first Hidden layer (Figure 9a,c), the other (second and third but not the first) Hidden layer(s) represent the dissimilarity between tasks better (Figure 9b,d,e).

**Figure 9.**
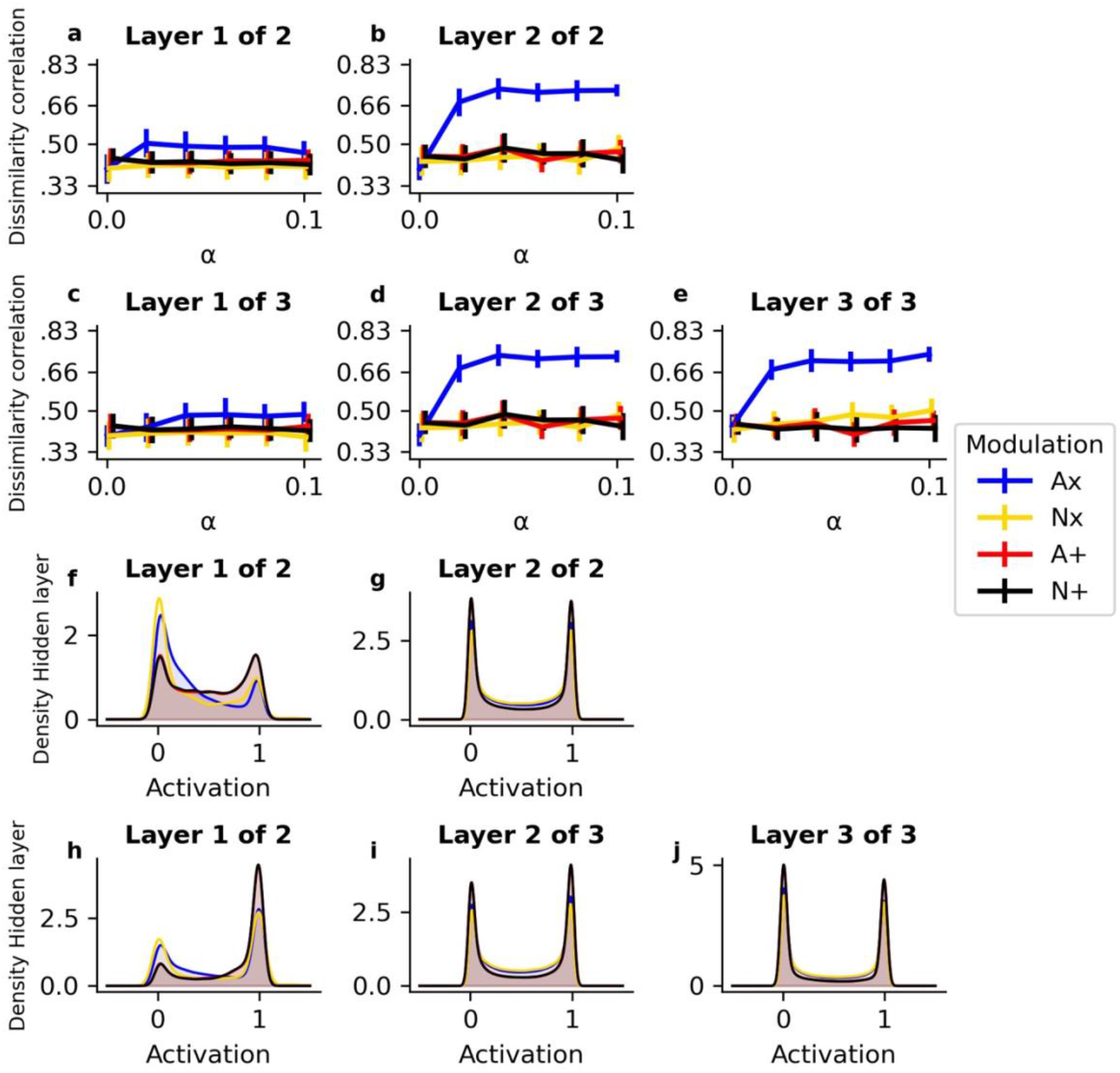
Hidden layer activity for multilayer networks. The upper two rows (panels a-e) illustrate the mean task dissimilarity correlation for each value of α across all simulations. Bars indicate 95% confidence intervals across 25 simulations. This is shown separately for each Hidden layer of the two Hidden layer network in (a-b) and for the three Hidden layer network (c-e). The lower two rows (f-j) illustrate the distribution of activation at each Hidden layer of the two-layer network (f-g) and the three-layer network (h- j).

Intriguingly, we observe in Figure 9f-j that, in terms of activation, the differences between the modulation networks are more pronounced at the first layer than at the second or third layer. This contrasts with the previous result (Figure 9a-e) that task dissimilarity correlations are more pronounced in the second and/or third Hidden layer(s). However, it is important to keep in mind that separating task representations does not necessarily lead to higher dissimilarity correlations as the different tasks also illustrate significant similarities. This is also emphasized by the fact that the Nx network clearly separates all tasks but does not show a strong dissimilarity correlation. Although it deserves further investigation, it might well be that task mappings are maximally separated in the first Hidden layer and then (compositionally) recombined in the deeper layers.

### 3.5. The location of modulation

To gain insight in how network performance is influenced by the location of modulation we performed additional simulations of the two Hidden layer network on the MNIST dataset. Here, we explored performance when the network received no modulatory input from the Task layer, when modulation was applied at the first Hidden layer (as before), at the second Hidden layer, or at both Hidden layers. As can be observed in Figure 10a, all networks perform at chance level when no modulation is applied. When modulation is applied at one or more Hidden layers, the multiplicative modulating networks, and in particular the Ax network, outperforms the additive networks. Although network performance seems more reliable (narrow confidence intervals) with modulation at deeper and/or more Hidden layers, the increase in mean accuracy is very small.

**Figure 10.**
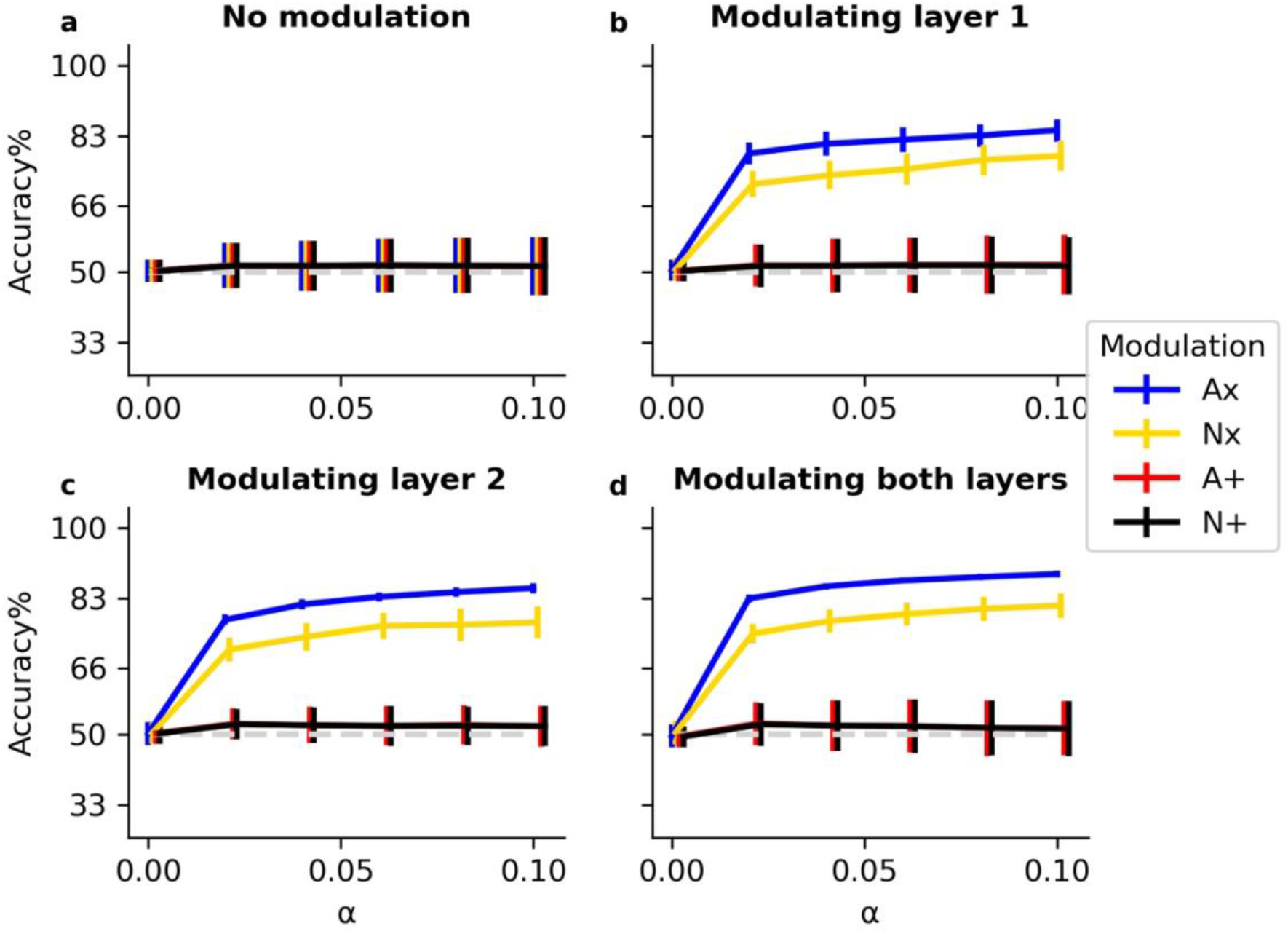
Accuracy in test phase for different locations of modulation. Lines illustrate mean accuracy during the test phase for each value of α, averaged across all tasks and all simulations. Bars indicate 95% confidence intervals over 25 simulations. The dashed lightgrey line indicates chance level accuracy. Results are shown for different datasets (rows) and different shapes of Hidden layer (columns).

**Figure 11.**
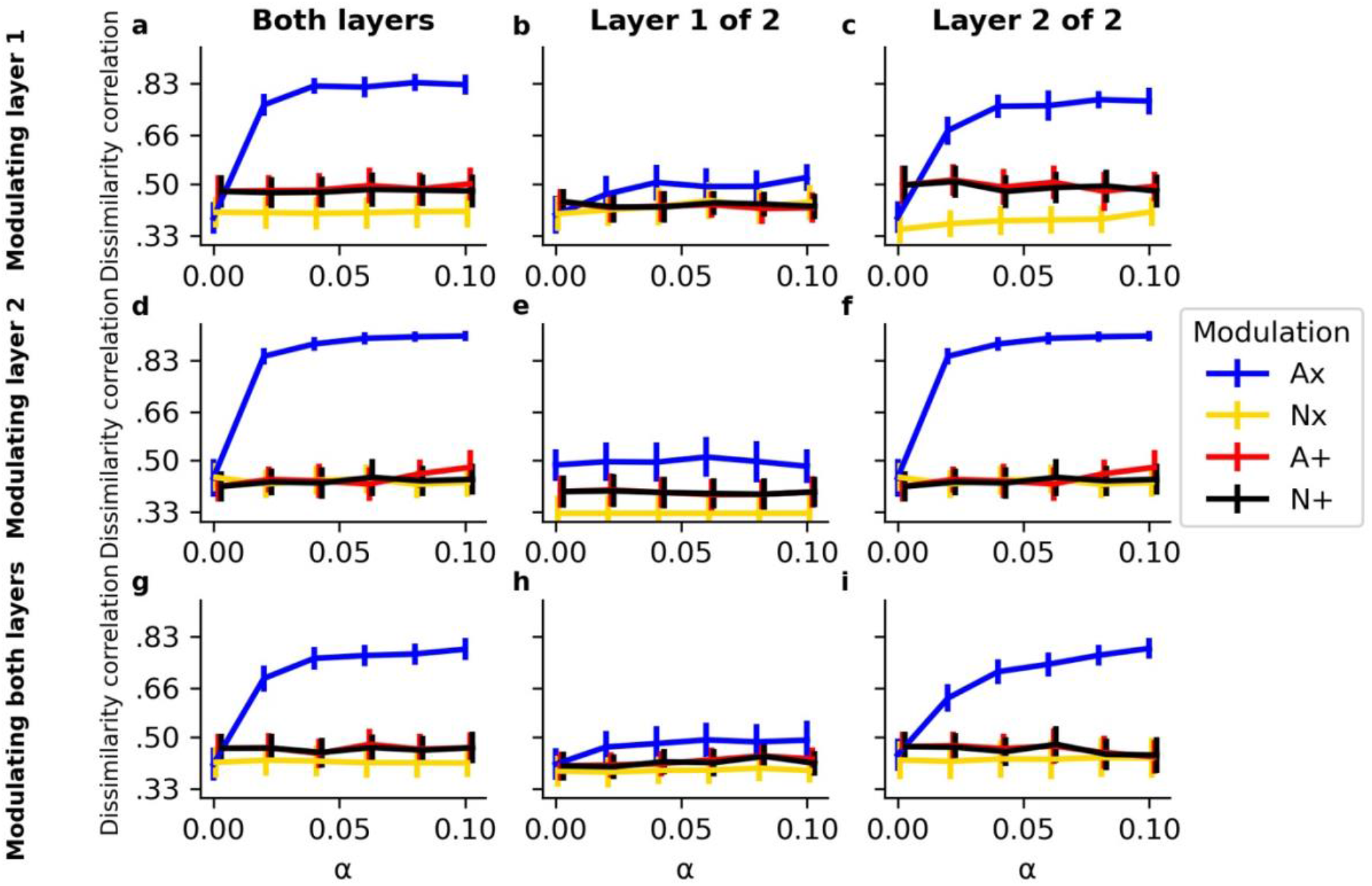
Correlation of objective and representational dissimilarity depending on location of modulation. Lines illustrate the mean task dissimilarity correlation for each value of α across all simulations. Bars indicate 95% confidence intervals over 25 simulations. Results are shown for different locations of modulation (rows) and different Hidden layer(s) (columns).

Also the results of task representational dissimilarity correlations seem highly similar for all locations of modulation. Note that this analysis could not be performed for the network that used no modulation. Since there was no task modulation, task representations were completely similar and the representational dissimilarity matrices for these networks were constant. Hence, no correlation could be computed with the objective dissimilarity matrix.

In sum, there does not seem to be a significant difference in network performance depending on where modulation is applied. In this specific case, modulation at layer 2 could be considered as most optimal since it requires to only learn 100 weights from each Task neuron to that second layer, compared to 300 for the first layer, and 400 for both layers.

## 4. Discussion

Current work investigated how neural networks can optimally balance the trade-off between avoiding interference via separating task representations and generalizing information via sharing representations. For this purpose, we identified and systematically investigated two crucial features of modulation signals. First, the modulation signals can be additive or multiplicative. Multiplicative signals were better suited for separating task representations. The multiplicative networks were less vulnerable to catastrophic interference than the additive networks. Second, the modulation signals could be adaptive (learned) or non-adaptive (random). Adaptive modulation signals provided a clear advantage over non-adaptive modulation signals in terms of both accuracy and balancing representations. Hence, the adaptive multiplicative (Ax) network was able to optimally balance the trade-off between sharing and separating task representations. This Ax network can avoid interference but also generalize across tasks which resulted in an overall better accuracy compared to the other networks.

Crucially, multiplicative signals modulated task-specific input more strongly. A Hidden neuron that receives input from many bottom-up Stimulus neurons, needs a strong (negative) additive modulation signal in order to be inhibited. In contrast, our multiplicative signal followed a RELU activation function (Equation (4); see Supplementary materials for an investigation of activation functions), which means that a small negative weight was sufficient to shut down (multiply by zero) a Hidden neuron activation. As a result, multiplicative signals developed sparser representations (Figure 7) which is optimal for separating task representations. This advantage was especially present when there were many Stimulus neurons (i.e., the MNIST task). When there were only 2 Stimulus neurons, as in the Trees task (see Figure 1c), the additive network was also able to separate task representations (see Figure 8h,k). Thus, multiplicative modulation is more efficient than additive signals, especially for separating high-dimensional inputs. Specifically, the Ax network warped the representational space in order to effectively organize tasks that obey similar mappings as well as tasks that obey dissimilar mappings within one neural architecture. In this representational space, dissimilar tasks were placed at the edges of a regular grid (see Figure 8c) and tasks that were similar were placed closer together. Such a geometrical organization of task rules is optimally suited for generalizing task rules (Bernardi et al., 2020; Kim, Pitt, & Myung, 2013). Consistent with the current analysis, Kim et al., (2013) demonstrated how backpropagation shapes hidden space to accommodate quasi-regularities in language processing, thus to accommodate both regular and irregular stimuli (e.g., orthography-phonology mappings). They demonstrated that after training by backpropagation, both regular and irregular stimuli could be placed in hidden neuron space at the edges of a slightly deformed grid; sufficiently grid-like to process the regular stimuli (and profit from generalization), but sufficiently deformed to cope with irregular mappings as well. Moreover, previous work has illustrated a similar systematicity of task representations in neural activation of prefrontal areas and hippocampus of monkeys (Bernardi et al., 2020).

An extensive amount of work describes how humans share or separate representations by extracting latent states in the environment (Collins & Frank, 2016; Franklin & Frank, 2018; Gershman & Niv, 2012; Wilson, Takahashi, Schoenbaum, & Niv, 2014; Yu, Wilson, & Nassar, 2020). Here, the agent decides on every new experience whether to categorize it as belonging to a new state or as belonging to a state that it has experienced before. Each latent state would then develop its own representations. An important disadvantage of the latent state approach is that it uses a dichotomous decision on whether an object belongs to the state or not. Such an approach is less suited to capture partial overlap between tasks. In the example of the Stroop dataset, a latent state approach would correctly assign tasks A, B and C to three different states because the mappings are completely dissimilar. The latent state approach would also correctly assign task E to the same latent state as A because they fully share the stimulus-action mappings. However, task D shares 1/3 of the mappings with all four other tasks. In this case, D would be optimally handled as a combination of the other mappings that are already learned. This is problematic for a latent state approach since it can only decide to assign D to a new latent state or to one of the previous ones.

To accommodate this limitation of the latent state approach, previous work has proposed compositionality (Fidler et al., 2009; Franklin & Frank, 2018; Lake et al., 2014; Sugita et al., 2011; Tubiana & Monasson, 2017; Yang et al., 2019). Specifically, the latent state approach could allow mixed overlap between tasks by representing a task as multiple states, each representing a subset of mappings (Franklin & Frank, 2020; Griffiths & Ghahramani, 2011). Nevertheless, this approach could significantly increase the number of possible states, which can be problematic in very complex task environments. This raises the question in how many states/dimensions the agent should cluster its experiences in order to optimally balance generalization and interference (Badre, Bhandari, Keglovits, & Kikumoto, 2021). As we have shown in Figure 4a,b, the current Ax network could benefit equally from the partial overlap in D and the full overlap between A and E without the need of extracting latent states. This is consistent with previous work, showing that multiplicative network interactions lead to useful compositional task representations (İrsoy & Cardie, 2015; Sugita et al., 2011).

Multiplicative modulation is also sometimes called gating (Masse et al., 2018; O’Reilly & Frank, 2006; Rougier, Noelle, Braver, Cohen, & O’Reilly, 2005). A crucial question that remains is how multiplicative signals are mechanistically implemented in the human brain. In this respect, we point to recent work that described an important role for neural oscillations in organizing functional networks. For example, it has been proposed that neural oscillations at alpha frequency (8-12 Hz) reflect gating by inhibition (Jensen & Mazaheri, 2010). Here, GABAergic inhibition provided by the task-irrelevant areas would be reflected by stronger alpha activity in those areas. Another oscillatory frequency that is known to organize functional networks in the brain is the theta frequency (4-8 Hz). More specifically, recent theoretical and empirical work (Helfrich & Knight, 2016; Lisman & Jensen, 2013; Verbeke et al., 2021; Verbeke & Verguts, 2019; Verguts, 2017) has proposed that prefrontal theta activity functions to (de)synchronize gamma (>40 Hz) activity in posterior processing areas. Here, synchronization leads to effective communication (gates open) between processing areas while desynchronization eliminates effective communication (gates closed) between processing areas (Fries, 2005, 2015). Thus, previous work has described how oscillatory interactions within and between different frequency bands (theta, alpha and gamma) reflect gating processes that could biologically implement the multiplicative modulation signals that were used in the current networks.

Previous work has also described how neurotransmitters such as dopamine and noradrenaline can modulate neural activation in a way that mimics multiplicative modulation (O’Reilly & Frank, 2006; Servan-Schreiber, Printz, & Cohen, 1990). Moreover, current work observed that additive modulation can reach similar performance as multiplicative modulation when stimulus dimensionality was low (Trees dataset). Hence, combining additive modulation with weight regularization between Stimulus and Hidden layer but not between Task and Hidden layer could allow the additive modulation networks to overcome high-dimensional inputs and reach similar performance as the multiplicative modulation networks. Future work should further explore biologically plausible implementations of multiplicative modulation.

In analogy to previous work (e.g., Cohen et al., 1990), the current networks used a low-dimensional Task layer which sent modulation signals to modulate a higher-dimensional Hidden layer. Typically, information in the Task layer has been considered to correspond to dorsolateral prefrontal cortex (DLPFC) and the Hidden layer to posterior task-related (e.g., visual and motor) processing pathways (Miller & Cohen, 2001). Hence, we propose that modulation signals are implemented by DLPFC.

A detailed investigation of activation at Hidden layer (Figure 7) illustrated that (multiplicative) networks separate tasks by developing sparse representations (see also Bowers, Vankov, Damian, & Davis, 2014). Here, for every task, only a small subset of neurons is active. In the current context, this suggests that the network develops groups or modules of neurons that each become specialized for a given task. Developing specialized modules might be beneficial for cognition in various ways (Bullinaria, 2007; Clune, Mouret, & Lipson, 2013; Coltheart, 1999; Fodor, 1983; Meunier, Lambiotte, Fornito, Ersche, & Bullmore, 2009). Moreover, recent work in reinforcement learning has also described hierarchical forms of modularity (Botvinick, Niv, & Barto, 2009; Dietterich, 2000; Holroyd & Verguts, 2021; Krueger & Dayan, 2009). Here, it is proposed that the dorsal part of the anterior cingulate cortex processes prediction errors related to specific events while the rostral part processes prediction errors related to the context in which these events occur (Alexander & Brown, 2015). However, modularity of processes in a single task requires integration of information across stages of processing. Hence, exploring the trade-off between sharing and separating representations at different levels of processing is an important avenue for future research.

In the current work, modulation signals are employed to guide learning over trials. Here, (for the adaptive networks) weights between the Task and Hidden layer are adapted to learn representations of different tasks. In contrast, a lot of previous work considered how modulation is adapted to guide online performance. Here, the intensity of the modulation signal is typically increased in response to some evaluation of the cost-benefit structure of the task context (Shenhav, Botvinick, & Cohen, 2013). These networks typically adapt the intensity of the modulation signal by changing activity in the Task layer instead of changing the weights (Botvinick et al., 2001; Verbeke & Verguts, 2020; Verguts, 2017). Consistent with this approach, research on visual attention has proposed that activity can be modulated (in a multiplicative manner) at the level of the Stimulus layer (Martinez-Trujillo & Treue, 2004; Treue & Martínez Trujillo, 1999). Alternatively, previous work (Cheadle et al., 2014) suggested that decisions can be guided via adaptive (multiplicative) gain functions in the transfer from input to output. Hence, there is a potentially important functional distinction between modulation in learning versus performance (see also Lindsay & Miller, 2017).

The trade-off between shared and separated representation also impacts performance (Musslick et al., 2020) Specifically, a wide range of empirical observations of interference during multi- tasking (performing multiple tasks at the same time) can be explained by a tendency to share task representations. Musslick et al. (2020) suggest that for generalization purposes, representations of novel tasks should strongly overlap with other task representations; unfortunately, such overlap leads to strong interference when these tasks need to be performed at the same time. However, with extensive training, the networks will gradually separate task representations, which leads to less interference. Hence, future work should consider a more extensive exploration of modulation signals in performance as well.

We evaluated the ability of different modulation signals to balance shared and separated representations by investigating how the networks could overcome catastrophic interference. Importantly, while modulation has proven to be efficient (Masse et al., 2018; Verbeke & Verguts, 2019), previous work also described other methods to avoid catastrophic interference. For instance, sharing or separation of task sets might be implemented in complementary learning systems (O’Reilly & Norman, 2002). Alternatively, machine learning literature has introduced methods such as synaptic intelligence (Kirkpatrick et al., 2017) in which weights learn whether they should be specific for one task and hence not change during further learning or whether they can be shared across all tasks. Future work should further address the differences between these approaches.

Several additional tests and extensions can be made to the network. First, previous work (Flesch et al., 2018) has pointed to an important distinction between interference in artificial and human agents. While artificial agents show more interference when they learn in a blocked fashion, human agents exhibit more interference when they learn in an interleaved fashion. The current work trained artificial agents in a blocked fashion. However, the different types of modulation signals should also be evaluated for interleaved learning. Potentially this approach could yield more insight in the existing distinction of learning benefits for artificial compared to human agents. Second, the current networks did not learn which task features were relevant for modulation. Here, the input layer was divided a-priori in a Stimulus group and a Task group. Hence, an important next step for the network would be to learn a hierarchical structure in input features in order to extract which inputs are relevant for modulation and which are relevant for basic stimulus-action mappings (Rougier et al., 2005). Alternatively, previous work has illustrated that compositional representations can even develop without providing task input if they are considered useful as situational signals (Butz et al., 2021). Third, although we illustrated that the Ax network was able to significantly benefit (in terms of accuracy) from shared mappings between tasks (Figure 4a,b), we did not perform a direct test of generalization. That is, we did not evaluate whether newly learned mappings in Task A of the Stroop task were transferred to task E without further training in task E. That is, we did not test whether the learned task relations were also suited for few-shot learning (Lake et al., 2014; Sylvain, Petrini, & Hjelm, 2020). Note that for an exact transfer between A and E, the weight matrix between Task neuron A and the Hidden layer should be exactly the same as the weight matrix between Task neuron E and the Hidden layer, which seems a strong requirement. Yet, it is not clear either whether two contexts that require the same stimulus-action mappings would also function in exactly the same way at behavioral and neural levels. Nor is this computationally desirable: In the natural environment, two labels that lead to the same stimulus-action contingencies, may still suggest (subtle) differences. Consider looking for lunch and seeing a “restaurant” versus a “snack bar” in the distance: Both tell you that you will be able to eat there, but expectations will differ at least slightly. As we discussed before, extracting a relational structure between contexts that allows for partial generalization might be more optimal than using a dichotomous same or different decision.

In sum, efficient human learning and performance requires to balance a trade-off between sharing representations to allow generalization and separating representations to avoid interference. We evaluated four different modulation signals and found that an adaptive multiplicative modulation signal was best suited to balance the sharing/separation trade-off. This modulation signal allowed the Hidden layer of the network to make a geometrical abstraction of the relational structure between tasks. Importantly, our work opens several avenues for future work to increase the understanding of the sharing/separation trade-off in both artificial and human agents.

## Acknowledgements

We thank Sebastian Musslick, Senne Braem, Cristian Buc Calderon and Pieter Huycke for valuable comments on this work. We also thank Michael J. Frank and his lab members for interesting input.

## Supplementary materials

### S.1. Exploration of activation functions

In the main text (see Equation (2)), multiplicative modulation was established by combining two nonlinear transformations of the Stimulus and Task input via *f*() and *g*(). Here, *f*() represented the sigmoid activation function (see Equation (3)) and *g*() represented a RELU activation function (see Equation (4)). Additive modulation (see Equation (1)) was implemented by transforming both the Task and Stimulus input with *f*(). Here, we present 4 novel simulations in which we explored all combinations of activation functions. Specifically, we tested networks in which both *f*() and *g*() corresponded to the sigmoid function (i.e. Sig (⊗ Sig)), we tested when *f*() corresponded to the sigmoid function and *g*() to the RELU function (i.e. Sig (⊗ RELU); as in the main text), we tested when *f*() corresponded to the RELU function and *g*() to the sigmoid function (i.e. RELU (⊗ Sig)) and we tested the networks when both *f*() and *g*() corresponded to the RELU function (i.e. RELU (⊗ RELU)). Note that for the additive modulation networks, only *f*() is relevant. This is why our notation shows the second function (i.e., *g*()) between brackets.

For these additional simulations, the networks were tested on the Stroop and Trees dataset with 12 Hidden neurons. We again explored different values of α ranging from 0 to 1 in steps of .2. Again, 25 simulations were performed in which we shuffled task contexts and trained networks for three context repetitions after which weights were frozen and the networks were tested again on each context.

Accuracy and dissimilarity correlations were analysed during the test phase. As can be observed (Figure S1), the network that is presented in the main text Sig (⊗ RELU) is most optimal for both the multiplicative and additive modulation networks. Although the multiplicative modulation seemed less efficient when implemented via a sigmoid function (Figure S1a,c,e,g,i,k,m,o), the Ax network still outperformed the other modulation methods. Presumably, the RELU function is more efficient for modulation because it operates at a much larger scale [0, +∞] than the sigmoid function which has bounds at 0 and 1.

**Figure S1.**
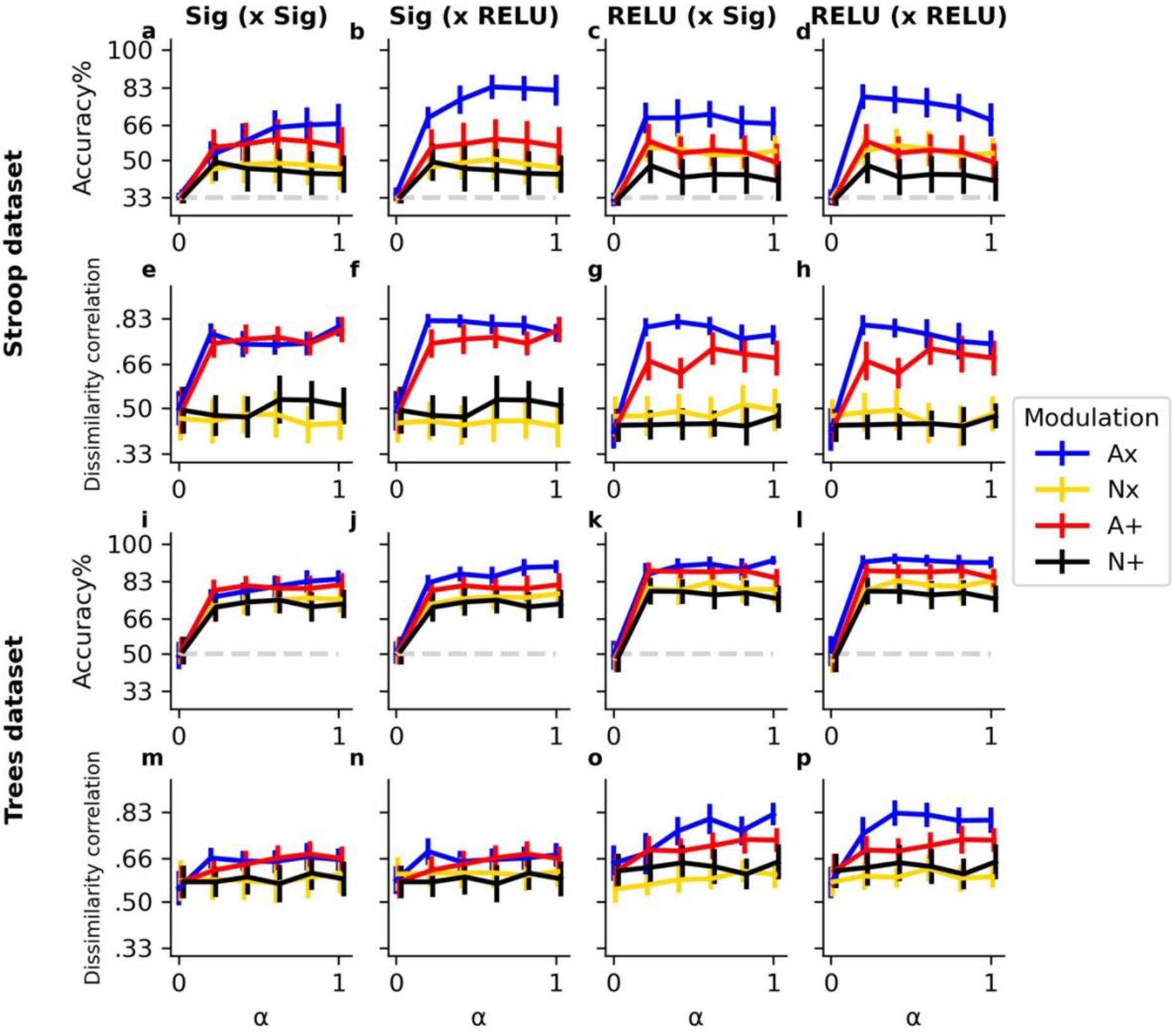
Activation function exploration. Lines illustrate mean accuracy/dissimilarity correlations for each value of α across all tasks and all simulations. Bars indicate 95% confidence intervals over 25 simulations. The dashed lightgrey line indicates chance level accuracy. Results are shown for different datasets (rows) and different combinations of activation functions (columns).

### S.2. Concatenated versus separated input transformations

In the main text (see Equation (2)), multiplicative modulation was established by combining two separated nonlinear transformations of the Stimulus and Task input via *f(SW)*⊗ *g(TW)* while Additive modulation (see Equation (1)) was implemented by transforming both the Task and Stimulus input with *f(SW+TW)*. Hence, for additive modulation, inputs were concatenated in one transformation, while for multiplicative modulation inputs were separated in two transformations. We choose these different implementations because they are most closely related to previous work (e.g., Cohen, Dunbar, & McClelland, 1990; Masse, Grant, & Freedman, 2018 for additive and multiplicative modulation respectively). However, for completeness, we also explored other implementations of multiplicative and additive modulation. Specifically, we tested networks in which for both types of modulation (additive and multiplicative) the inputs were concatenated. This results in *f*(*SW*) ⊗ *g*(*TW*) for multiplicative modulation and *f*(*SW*) + *g*(*TW*) for additive modulation. Additionally, we tested networks in which the inputs were separated. This resulted in *f*(*SW* ⊗ *TW*) for multiplicative modulation and *f*(*SW* + *TW*) for additive modulation.

Again, the networks were tested on the Stroop and Trees dataset with 12 Hidden neurons. We explored different values of α going from 0 to 1 in steps of .2 and 25 simulations were performed in which we shuffled task contexts and trained networks for three context repetitions after which weights were frozen and the networks were tested again on each context.

Accuracy and dissimilarity correlations were analysed during the test phase. As illustrated in (Figure S2), for none of the modulation methods there is a clear difference when using separated versus concatenated input transformations.

**Figure S2.**
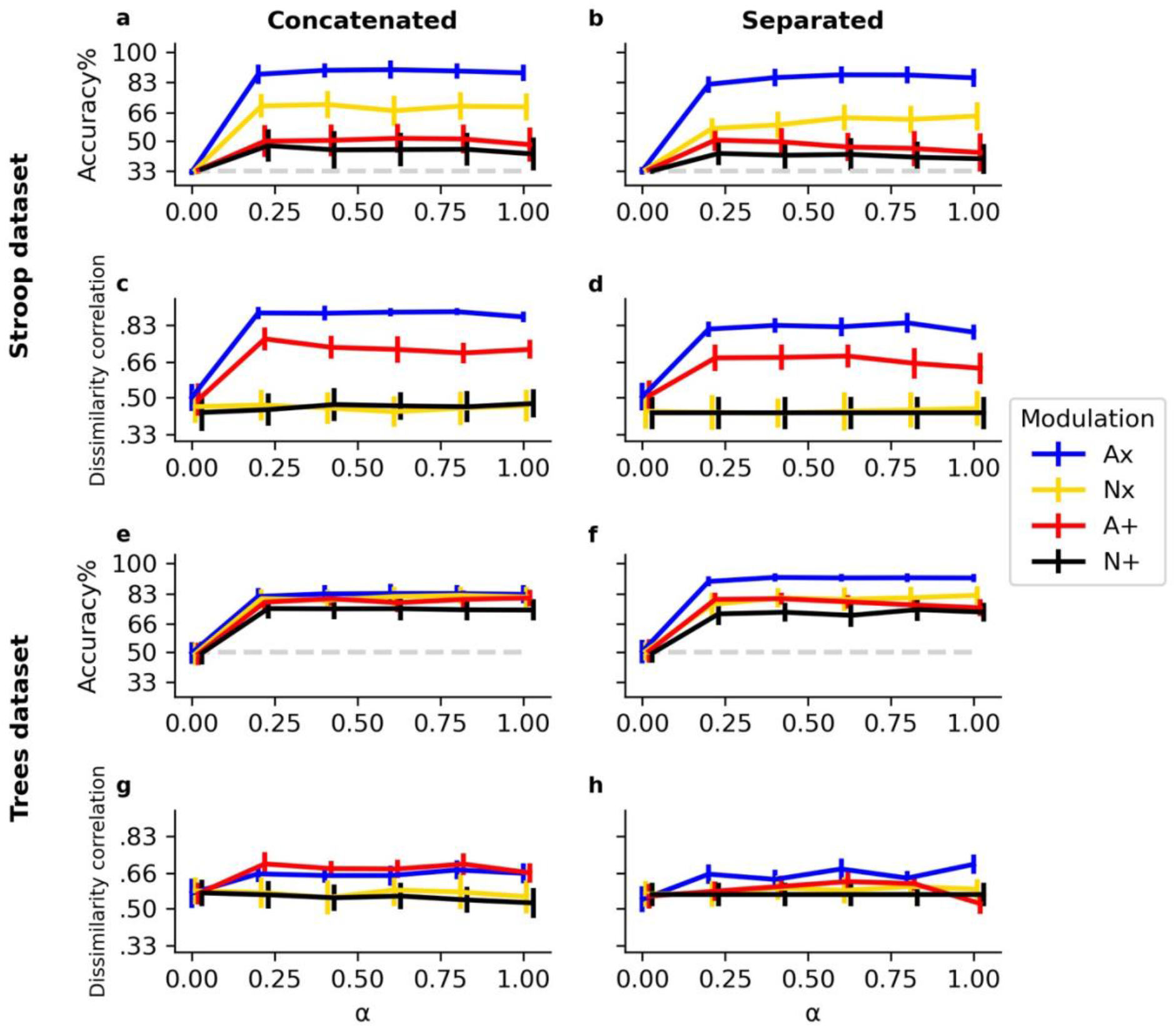
Concatenated versus separated input transformations. Lines illustrate mean accuracy/dissimilarity correlations for each value of α across all tasks and all simulations. Bars indicate 95% confidence intervals over 25 simulations. The dashed lightgrey line indicates chance level accuracy. Results are shown for different datasets (rows) and different concatenated versus separated input transformations (columns).

### S.3. Exploration of weight initialization

In the main text we described that all weights are initialized with a random value drawn from the normal distribution N(0, 1). Only for the Ax network, modulating weights (between Task and Hidden layer) had an initial random value drawn from the uniform distribution U(0, 1), such that RELU(***T***) > 0 (all gates open) at the first trial. However, previous work has demonstrated that the way in which weights are initialized is not trivial (e.g., Flesch, Juechems, Dumbalska, & Saxe, 2021). To illustrate that our results were not solely driven by this choice of weight initialization, we performed additional simulations in which we tested a normal initialization for all modulating weights and compared this to a uniform initialization for all modulating weights.

The networks were again tested on the Stroop and Trees dataset with 12 Hidden neurons. We explored different values of α ranging from 0 to 1 in steps of .2. Again, 25 simulations were performed in which we shuffled task contexts and trained networks for three context repetitions after which weights were frozen and the networks were tested again on each context.

As shown in Figure S3, the Ax network indeed performs a bit worse when the weights are initialized from the normal distribution. However, it is also the case that for the other three networks the uniform initialization was a bit less optimal. Hence, the main text describes simulations in which the optimal weight initialization was chosen for each of the four modulation methods.

**Figure S2.**
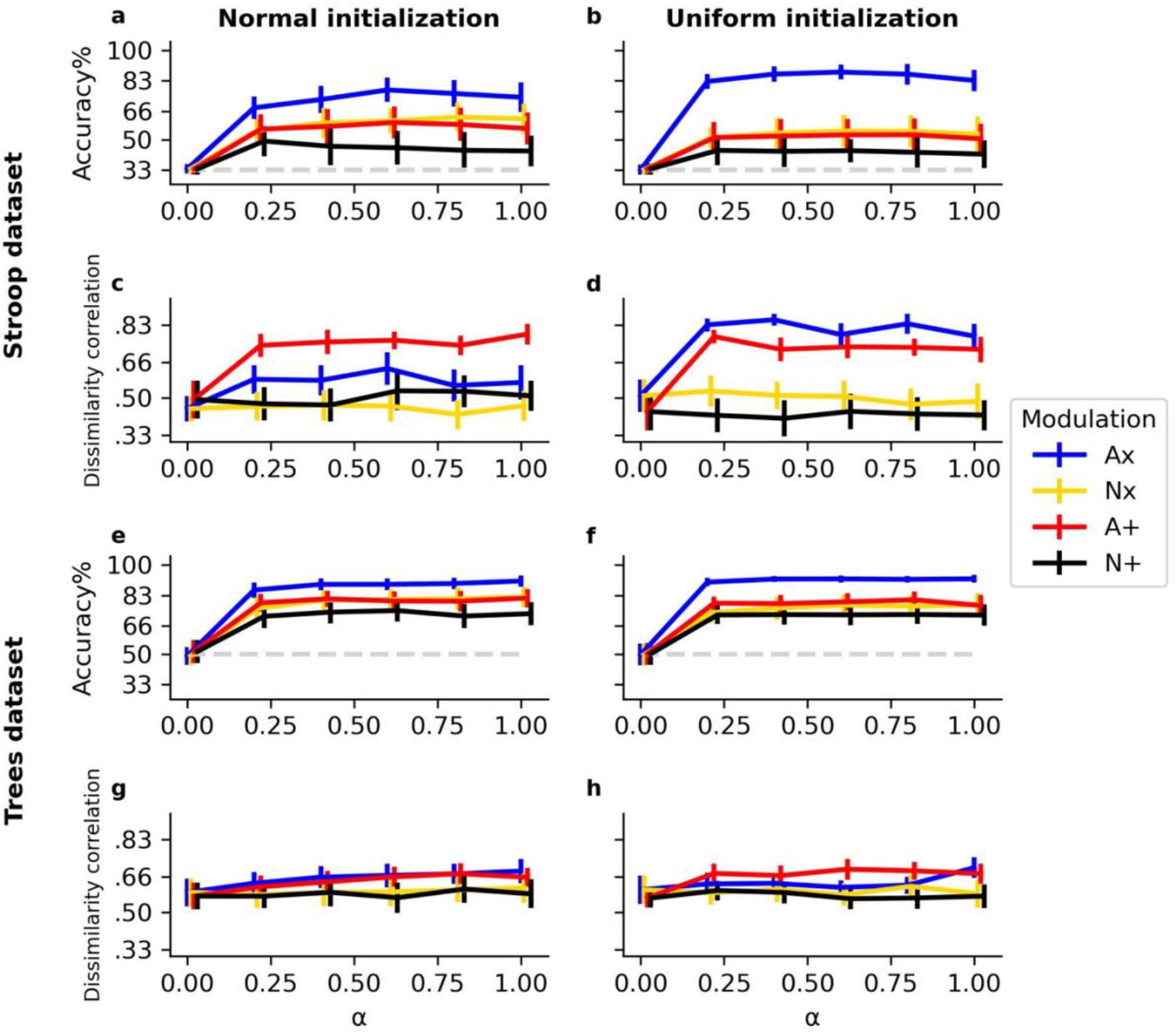
Weight initialization exploration. Lines illustrate mean accuracy/dissimilarity correlations for each value of α across all tasks and all simulations. Bars indicate 95% confidence intervals over 25 simulations. The dashed lightgrey line indicates chance level accuracy. Results are shown for different datasets (rows) and for different initializations of the modulation weights (columns).

## References

1. Aben, B., Calderon, C. B., Van den Bussche, E., & Verguts, T. (2020). Cognitive Effort Modulates Connectivity between Dorsal Anterior Cingulate Cortex and Task-Relevant Cortical Areas. The Journal of Neuroscience, 40(19). https://doi.org/10.1523/jneurosci.2948-19.2020

2. Abrahamse, E., Braem, S., Notebaert, W., & Verguts, T. (2016). Grounding Cognitive Control in Associative Learning. Psychological Bulletin, 142(7), 693–728. https://doi.org/10.1037/bul0000047

3. Alexander, W. H., & Brown, J. W. (2015). Hierarchical error representation: A computational model of anterior cingulate and dorsolateral prefrontal cortex. Neural Computation, 27(11), 2354– 2410. https://doi.org/10.1162/NECO

4. Alon, N., Reichman, D., Shinkar, I., Wagner, T., Musslick, S., Cohen, J. D., … Ozcimder, K. (2017). A graph-theoretic approach to multitasking. In Advances in Neural Information Processing Systems (pp. 2097–2106).

5. Badre, D., Bhandari, A., Keglovits, H., & Kikumoto, A. (2021). The dimensionality of neural representations for control. Current Opinion in Behavioral Sciences, 38, 20–28. https://doi.org/10.1016/j.cobeha.2020.07.002

6. Baxter, J. (2019). Learning Internal Representations. In Proceedings of the eigth annual conference on computational learning theory (pp. 311–320). https://doi.org/10.7551/mitpress/2906.003.0006

7. Bernardi, S., Benna, M. K., Rigotti, M., Munuera, J., Fusi, S., & Salzman, C. D. (2020). The Geometry of Abstraction in the Hippocampus and Prefrontal Cortex. Cell, 183(4), 954–967. https://doi.org/10.1016/j.cell.2020.09.031

8. Botvinick, M. M., Braver, T. S., Barch, D. M., Carter, C. S., & Cohen, J. D. (2001). Conflict monitoring and cognitive control. Psychological Review, 108(3), 624–652. https://doi.org/10.1037/0033-295X.108.3.624

9. Botvinick, M. M., Niv, Y., & Barto, A. C. (2009). Hierarchically organized behavior and its neural foundations : A reinforcement learning perspective. Cognition, 113(3), 262–280. https://doi.org/10.1016/j.cognition.2008.08.011

10. Bouchacourt, F., & Buschman, T. J. (2019). A Flexible Model of Working Memory. Neuron, 103(1), 147–160. https://doi.org/10.1016/j.neuron.2019.04.020

11. Bowers, J. S., Vankov, I. I., Damian, M. F., & Davis, C. J. (2014). Neural networks learn highly selective representations in order to overcome the superposition catastrophe. Psychological Review, 121(2), 248–261. https://doi.org/10.1037/a0035943

12. Bullinaria, J. A. (2007). Understanding the advantage of modularity in neural systems. Cognitive Science, 31(4), 673–695. https://doi.org/10.1080/15326900701399939

13. Butz, M. V., Achimova, A., Bilkey, D., & Knott, A. (2021). Event-Predictive Cognition: A Root for Conceptual Human Thought. Topics in Cognitive Science, 13(1), 10–24. https://doi.org/10.1111/tops.12522

14. Butz, M. V., Bilkey, D., Humaidan, D., Knott, A., & Otte, S. (2019). Learning, planning, and control in a monolithic neural event inference architecture. Neural Networks, 117, 135–144. https://doi.org/10.1016/j.neunet.2019.05.001

15. Cheadle, S., Wyart, V., Tsetsos, K., Myers, N., deGardelle, V., HerceCastañón, S., & Summerfield, C. (2014). Adaptive gain control during human perceptual choice. Neuron, 81(6), 1429–1441. https://doi.org/10.1016/j.neuron.2014.01.020

16. Clune, J., Mouret, J. B., & Lipson, H. (2013). The evolutionary origins of modularity. Proceedings of the Royal Society B: Biological Sciences, 280(1755). https://doi.org/10.1098/rspb.2012.2863

17. Cohen, J. D., Dunbar, K., & McClelland, J. L. (1990). On the control of automatic processes: a parallel distributed processing account of the Stroop effect. Psychological Review, 97(3), 332–361. https://doi.org/10.1037/0033-295X.97.3.332

18. Collins, A. G. E., & Frank, M. J. (2016). Neural signature of hierarchically structured expectations predicts clustering and transfer of rule sets in reinforcement learning. Cognition, 152, 160–169. https://doi.org/10.1016/j.cognition.2016.04.002

19. Coltheart, M. (1999). Modularity and cognition. Trends in Cognitive Sciences, 3(3), 115–120. https://doi.org/10.1016/S1364-6613(99)01289-9

20. Dietterich, T. G. (2000). Hierarchical reinforcement learning with the MAXQ value function decomposition. Journal of Artificial Intelligence Research, 13, 227–303. https://doi.org/10.1613/jair.639

21. Fidler, S., Boben, M., & Leonardis, A. (2009). Learning hierarchical compositional representations of object structure. Object Categorization: Computer and Human Vision Perspectives, 9780521887, 196–215. https://doi.org/10.1017/CBO9780511635465.012

22. Flesch, T., Balaguer, J., Dekker, R., Nili, H., & Summerfield, C. (2018). Comparing continual task learning in minds and machines. Proceedings of the National Academy of Sciences of the United States of America, 115(44). https://doi.org/10.1073/pnas.1800755115

23. Fodor, J. A. (1983). The modularity of mind. MIT Press/Bradford Books.

24. Franklin, N. T., & Frank, M. J. (2018). Compositional clustering in task structure learning. PLoS Computational Biology, 14(4), 1–25. https://doi.org/10.1371/journal.pcbi.1006116

25. Franklin, N. T., & Frank, M. J. (2020). Generalizing to generalize: Humans flexibly switch between compositional and conjunctive structures during reinforcement learning. PLoS Computational Biology, 16(4), 1–33. https://doi.org/10.1371/journal.pcbi.1007720

26. French, R. M. (1999). Catastrophic forgetting in connectionist networks. Trends in Cognitive Sciences, 3(4), 128–135.

27. Fries, P. (2005). A mechanism for cognitive dynamics: neuronal communication through neuronal coherence. Trends in Cognitive Sciences, 9(10), 474–480. https://doi.org/10.1016/j.tics.2005.08.011

28. Fries, P. (2015). Rhythms for Cognition: Communication through Coherence. Neuron, 88(1), 220–235. https://doi.org/10.1016/j.neuron.2015.09.034

29. Gershman, S. J., & Niv, Y. (2012). Exploring a latent cause theory of classical conditioning. Learning and Behavior, 40(3), 255–268. https://doi.org/10.3758/s13420-012-0080-8

30. Griffiths, T. L., & Ghahramani, Z. (2011). The Indian buffet process: An introduction and review. Journal of Machine Learning Research, 12, 1185–1224.

31. Helfrich, R. F., & Knight, R. T. (2016). Oscillatory Dynamics of Prefrontal Cognitive Control. Trends in Cognitive Sciences, 20(12), 916–930. https://doi.org/10.1016/j.tics.2016.09.007

32. Holroyd, C. B., & Verguts, T. (2021). The best laid plans: Computational principles of ACC. Trends in Cognitive Sciences, 25(4), 316–329. https://doi.org/10.1016/j.tics.2021.01.008

33. Hupkes, D., Dankers, V., Mul, M., & Bruni, E. (2020). Compositionality decomposed: How do neural networks generalise? Journal of Artificial Intelligence Research, 67, 757–795.

34. Irsoy, O., & Cardie, C. (2014). Deep recursive neural networks for compositionality in language. Advances in Neural Information Processing Systems, 3(January), 2096–2104.

35. İrsoy, O., & Cardie, C. (2015). Modeling compositionality with multiplicative recurrent neural networks. 3rd International Conference on Learning Representations, ICLR 2015 - Conference Track Proceedings, (2013), 1–10.

36. Jensen, O., & Mazaheri, A. (2010). Shaping functional architecture by oscillatory alpha activity: Gating by inhibition. Frontiers in Human Neuroscience, 4, 1–8. https://doi.org/10.3389/fnhum.2010.00186

37. Kim, W., Pitt, M. A., & Myung, J. I. (2013). How Do PDP Models Learn Quasiregularity? Psychological Review, 120(4), 903–916. https://doi.org/10.1037/a0034195

38. Kirkpatrick, J., Pascanu, R., Rabinowitz, N., Veness, J., Desjardins, G., Rusu, A. A., … Hadsell, R. (2017). Overcoming Catastrophic Forgetting in Neural Networks. Proceedings of the National Academy of Sciences, 114(13), 3521–3526. https://doi.org/10.1073/pnas.1611835114

39. Krueger, K. A., & Dayan, P. (2009). Flexible shaping: How learning in small steps helps. Cognition, 110(3), 380–394. https://doi.org/10.1016/j.cognition.2008.11.014

40. Lake, B., & Baroni, M. (2018). Generalization without systematicity: On the compositional skills of sequence-to-sequence recurrent networks. 35th International Conference on Machine Learning, ICML 2018, 7, 4487–4499.

41. Lake, B., Lee, C.-Y., Glass, J., Lake, B. M., Glass, J. R., & Tenenbaum, J. B. (2014). One-shot learning of generative speech concepts Publication Date One-shot learning of generative speech concepts. Proceedings of the Annual Meeting of the Cognitive Science Society, (36), 803–808. Retrieved from https://cloudfront.escholarship.org/dist/prd/content/qt3xf2n3vc/qt3xf2n3vc.pdf

42. Lake, B., Ullman, T., Tenenbaum, J., & Gershman, S. (2017). Building machines that learn and think like people. Behavioral and Brain Sciences, 40(2017). https://doi.org/10.1017/S0140525X16001837

43. LeCun, Y., Cortes, C., & Burges, C. J. (2010). MNIST handwritten digit database.

44. Lillicrap, T. P., Cownden, D., Tweed, D. B., & Akerman, C. J. (2016). Random synaptic feedback weights support error backpropagation for deep learning. Nature Communications, 7, 1–10. https://doi.org/10.1038/ncomms13276

45. Lindsay, G. W., & Miller, K. D. (2018). How biological attention mechanisms improve task performance in a large-scale visual system model. ELife, 7, 1–29. https://doi.org/10.7554/eLife.38105.001

46. Lisman, J. E., & Jensen, O. (2013). The Theta-Gamma Neural Code. Neuron, 77(6), 1002–1016. https://doi.org/10.1016/j.neuron.2013.03.007

47. Maass, W., Natschläger, T., & Markram, H. (2002). Real-time computing without stable states: A new framework for neural computation based on perturbations. Neural Computation, 14(11), 2531–2560. https://doi.org/10.1162/089976602760407955

48. Martinez-Trujillo, J. C., & Treue, S. (2004). Feature-Based Attention Increases the Selectivity of Population Responses in Primate Visual Cortex. Current Biology, 14, 744–751. https://doi.org/10.1016/j.cub.2004.04.028

49. Masse, N. Y., Grant, G. D., & Freedman, D. J. (2018). Alleviating catastrophic forgetting using context-dependent gating and synaptic stabilization. Proceedings of the National Academy of Sciences, 115(44), 1–12. https://doi.org/10.1073/pnas.1803839115

50. McClelland, J. L., McNaughton, B. L., & O’Reilly, R. C. (1995). Why There Are Complementary Learning Systems in the Hippocampus and Neo-cortex: Insights from the Successes and Failures of Connectionists Models of Learning and Memory. Psychological Review., 102(3), 419–457. https://doi.org/10.1037/0033-295X.102.3.419

51. Meunier, D., Lambiotte, R., Fornito, A., Ersche, K. D., & Bullmore, E. T. (2009). Hierarchical modularity in human brain functional networks. Frontiers in Neuroinformatics, 3, 1–12. https://doi.org/10.3389/neuro.11.037.2009

52. Miller, E. K., & Cohen, J. D. (2001). An Integrative Theory of Prefrontal Cortex Function. Annual Review of Neuroscience, 24, 167–202. https://doi.org/10.1146/annurev.neuro.24.1.167

53. Musslick, S., & Cohen, J. D. (2020). Rationalizing constraints on the capacity for cognitive control. PsyArxiv, 45. https://doi.org/10.31234/osf.io/vtknh

54. Musslick, S., Saxe, A. M., Ozcimder, K., Dey, B., Henselman, G., & Cohen, J. D. (2017). Multitasking Capability Versus Learning Efficiency in Neural Network Architectures. In Annual meeting of the Cognitive Science Society (pp. 829–834).

55. Musslick, S., Saxe, A., Novick, A., Reichman, D., & Cohen, J. D. (2020). On the rational boundedness of cognitive control: Shared versus separated representations. PsyArXiv. https://doi.org/10.31234/osf.io/jkhdf

56. O’Reilly, R. C., & Frank, M. J. (2006). Making Working Memory Work : A Computational Model of Learning in the Prefrontal Cortex and Basal Ganglia. Neural Computation, 18(2), 283–328. https://doi.org/10.1162/089976606775093909

57. O’Reilly, R. C., & Norman, K. A. (2002). Hippocampal and neocortical contributions to memory: Advances in the complementary learning systems framework. Trends in Cognitive Sciences, 6(12), 505–510. https://doi.org/10.1016/S1364-6613(02)02005-3

58. Rougier, N. P., Noelle, D. C., Braver, T. S., Cohen, J. D., & O’Reilly, R. C. (2005). Prefrontal cortex and flexible cognitive control: Rules without symbols. Proceedings of the National Academy of Sciences of the United States of America, 102(20), 7338–7343. https://doi.org/10.1073/pnas.0502455102

59. Rumelhart, D. E., Hinton, G. E., & Williams, R. J. (1986). Learning representations by back- propagating errors. Nature, 323, 533–536. https://doi.org/10.1038/323533a0

60. Sagiv, Y., Musslick, S., Niv, Y., & Cohen, J. D. (2020). Efficiency of learning vs. processing: Towards a normative theory of multitasking. In Proceedings of the 40th Annual Meeting of the Cognitive Science Society, (p. 1004—1009).

61. Servan-Schreiber, D., Printz, H., & Cohen, J. D. (1990). A network model of catecholamiine effects: Gain, signal-to-noise ratio, and behavior. Science, 249(4971), 892–895. https://doi.org/10.1126/science.2392679

62. Shenhav, A., Botvinick, M. M., & Cohen, J. D. (2013). The expected value of control: An integrative theory of anterior cingulate cortex function. Neuron, 79(2), 217–240. https://doi.org/10.1016/j.neuron.2013.07.007

63. Stroop, J. R. (1935). Studies of interference in serial verbal reactions. Journal of Experimental Psychology, 18(6), 643–662. https://doi.org/10.1037/h0054651

64. Sugita, Y., Tani, J., & Butz, M. V. (2011). Simultaneously emerging braitenberg codes and compositionality. Adaptive Behavior, 19(5), 295–316. https://doi.org/10.1177/1059712311416871

65. Sylvain, T., Petrini, L., & Hjelm, R. D. (2020). Zero-Shot Learning from scratch (ZFS): leveraging local compositional representations. Retrieved from http://arxiv.org/abs/2010.13320

66. Treue, S., & Martínez Trujillo, J. C. (1999). Feature-based attention influences motion processing gain in macaque visual cortex. Nature, 399, 575–579. https://doi.org/10.1038/21176

67. Tsai, C. Y., Saxe, A., & Cox, D. (2016). Tensor switching networks. Advances in Neural Information Processing Systems, (Nips), 2046–2054.

68. Tubiana, J., & Monasson, R. (2017). Emergence of Compositional Representations in Restricted Boltzmann Machines. Physical Review Letters, 118(13), 1–5. https://doi.org/10.1103/PhysRevLett.118.138301

69. Vaidya, A. R., Jones, H. M., Castillo, J., & Badre, D. (2021). Neural representation of abstract task structure during generalization. ELife, (10:e63226.), 1–22. https://doi.org/10.7554/eLife.63226

70. Verbeke, P., Ergo, K., De Loof, E., Verguts, T., Loof, E. De, & Verguts, T. (2021). Learning to synchronize: Midfrontal theta dynamics during rule switching. Journal of Neuroscience, 41(7), 1–13. https://doi.org/10.1523/JNEUROSCI.1874-20.2020

71. Verbeke, P., & Verguts, T. (2019). Learning to synchronize: How biological agents can couple neural task modules for dealing with the stability-plasticity dilemma. PLoS Computational Biology, 15(8). https://doi.org/10.1371/journal.pcbi.1006604

72. Verbeke, P., & Verguts, T. (2020). Neural synchrony for adaptive control. Psyarxiv, 1–37. https://doi.org/10.31234/osf.io/523x9

73. Verguts, T. (2017). Binding by random bursts: A computational model of cognitive control. Journal of Cognitive Neuroscience, 29(6), 1103–1118. https://doi.org/10.1162/jocn

74. Verguts, T., & Notebaert, W. (2008). Hebbian Learning of Cognitive Control : Dealing With Specific and Nonspecific Adaptation. Psychological Review, 115(2), 518–525. https://doi.org/10.1037/0033-295X.115.2.518

75. Wilson, R. C., Takahashi, Y. K., Schoenbaum, G., & Niv, Y. (2014). Orbitofrontal cortex as a cognitive map of task space. Neuron, 81(2), 267–279. https://doi.org/10.1016/j.neuron.2013.11.005

76. Yang, G. R., Joglekar, M. R., Song, H. F., Newsome, W. T., & Wang, X. J. (2019). Task representations in neural networks trained to perform many cognitive tasks. Nature Neuroscience, 22(2), 297–306. https://doi.org/10.1038/s41593-018-0310-2

77. Yu, L. Q., Wilson, R. C., & Nassar, M. R. (2020). Adaptive learning is structure learning in time. Psyarxiv, 1–27. https://doi.org/10.31234/osf.io/r637c

78. Zambaldi, V., Raposo, D., Santoro, A., Bapst, V., Li, Y., Babuschkin, I., … Battaglia, P. (2018). Relational Deep Reinforcement Learning, (2), 1–15. Retrieved from http://arxiv.org/abs/1806.01830

## References

1. Cohen, J. D., Dunbar, K., & McClelland, J. L. (1990). On the control of automatic processes: a parallel distributed processing account of the Stroop effect. Psychological Review, 97(3), 332–361. https://doi.org/10.1037/0033-295X.97.3.332

2. Flesch, T., Juechems, K., Dumbalska, T., & Saxe, A. (2021). Rich and lazy learning of task representations in brains and neural networks. BioRxiv.

3. Masse, N. Y., Grant, G. D., & Freedman, D. J. (2018). Alleviating catastrophic forgetting using context-dependent gating and synaptic stabilization. Proceedings of the National Academy of Sciences, 115(44), 1–12. https://doi.org/10.1073/pnas.1803839115

